# Decreased fucosylation impacts epithelial integrity and increases risk for COPD

**DOI:** 10.1101/2023.10.31.564805

**Authors:** Carter Swaby, Bonnie Yeung-Luk, Shreeti Thapa, Kristine Nishida, Arabelis Wally, Baishakhi Ghosh, Austin Niederkofler, Sean Luk, Mirit Girgis, Allison Keller, Cecilia Cortez, Sahana Ramaswamy, Kai Wilmsen, Laura Bouché, Anne Dell, M. Bradley Drummond, Nirupama Putcha, Stuart M. Haslam, Rasika Mathias, Nadia N. Hansel, Jian Sheng, Venkataramana Sidhaye

## Abstract

COPD causes significant morbidity and mortality worldwide. Epithelial damage is fundamental to disease pathogenesis, although the mechanisms driving disease remain undefined. Published evidence from a COPD cohort (SPIROMICS) and confirmed in a second cohort (COPDgene) demonstrate a polymorphism in *Fucosyltransferese-2 (FUT2)* is a trans-pQTL for E-cadherin, which is critical in COPD pathogenesis. We found by MALDI-TOF analysis that *FUT2* increased terminal fucosylation of E-cadherin. Using atomic force microscopy, we found that FUT2-dependent fucosylation enhanced E-cadherin-E-cadherin bond strength, mediating the improvement in monolayer integrity. Tracheal epithelial cells from *Fut2*^-/-^ mice have reduced epithelial integrity, which is recovered with reconstitution of *Fut2*. Overexpression of *FUT2* in COPD derived epithelia rescues barrier function. *Fut2^-/-^* mice show increased susceptibility in an elastase model of disease developing both emphysema and fibrosis. We propose this is due to the role of *FUT2* in proliferation and cell differentiation. Overexpression of FUT2 significantly increased proliferation. Loss of *Fut2* results in accumulation of Spc+ cells suggesting a failure of alveolar type 2 cells to undergo transdifferentiation to alveolar type 1. Using a combination of population data, genetically manipulated mouse models, and patient-derived cells, we present a novel mechanism by which post-translational modifications modulate tissue pathology and serve as a proof of concept for the development of a disease-modifying target in COPD.

## Introduction

Chronic obstructive pulmonary disease (COPD), characterized by airflow obstruction, emphysema, and epithelial barrier dysfunction, kills over three million people globally per year (1, 2). The primary cause of COPD in the US is long-term exposure to inhaled cigarette smoke (CS) (3) although other chronic insults such as air pollution significantly contribute to its global incidence(4). The lung epithelium is the first point of contact for inhalants and is responsible for serving as a barrier to prevent access to subepithelial tissues. E-cadherin, an adherens junction protein, regulates the permeability, polarization, and differentiation of the epithelium. As such, E-cadherin’s crucial role in the formation and maintenance of the lung epithelium is clear (5).

E-cadherin is most known for its role in maintaining calcium-dependent cell-cell adhesion in epithelial cells (6). However, studies have shown that it is involved in a wide range of cellular activities such as cell maturation, differentiation and migration, cell signaling, immune response, and tumor suppression (5, 7). This versatile role makes E-cadherin a protein of interest for numerous diseases, especially COPD. Decreases in E-cadherin in both the airways and the alveoli have long been associated with COPD (8–14). Our lab has demonstrated that primary bronchial epithelial cells derived from patients with COPD have a significant reduction in E-cadherin levels compared to age- and sex-matched normal cells (15, 16). Moreover, we have found that loss of E-cadherin can drive epithelial dysfunction and tissue remodeling (15) in mouse models. However, mechanisms of modulating E-cadherin in COPD are unknown.

Post-translational modifications are covalent changes to amino acids within a protein and can significantly alter protein function or stability. They are one of the last steps in protein biosynthesis and are independent of their original gene transcript. The ability of proteins to undergo post-translational modification at any stage allows for alterations to protein structure and function, which can have far-reaching effects on cellular function (17). One of the most common and highly regulated post-translational modifications is glycosylation, which plays a vital role in governing protein folding, stability, and protein-protein interactions (18). Glycosylation is based on enzymatic reactions that add glycans to proteins and encompass a wide selection of sugar moieties to specific amino acids. Glycosylation occurs in the endoplasmic reticulum and golgi apparatus (PMID 26956395, NCBI Gene 14344). Fucosylation, a specific type of glycosylation, is the process of transferring of a fucose sugar to its substrates, N- and O-linked glycans that are attached to a protein structure, by fucosyltransferases (FUTs) (19).

Fucosyltransferase-2 is an intracellular protein responsible for catalysis of α[1,2] fucosylation on the terminal galactose. This primarily occurs on glycan type 1 chain precursors with specificity for epithelial cells (20). Intracellularly, fucosyltransferase-2 is primarily localized in the Golgi apparatus. GDP-l-fucose is transported to the Golgi by a GDP-l-fucose transporter where it is then transferred to the glycan. Following their fucosylation and other modifications made in the Golgi apparatus, these fucosylated proteins are then shuttled to their final destinations in vesicles (21).

In this study, we combine clinical genomic data with purified protein analysis, genetically manipulated mouse models, patient-derived differentiated epithelia, and human precision-cut lung slices, to investigate the impact of terminal fucosylation of E-cadherin on epithelial integrity and susceptibility to lung damage from CS. We began with the identification of an SNP that results in loss of FUT2 that is associated with E-cadherin and COPD in two independent clinical cohorts. Having confirmed that FUT2 could post-translationally modify E-cadherin by mass spectrometry and immunoprecipitation, we studied mouse models and patient derived cells to assess its effect on monolayer integrity and lung morphometry.

## Results

### Co-localization of trans pQTL for E-cadherin and cis eQTL for FUT2

In a meta-analysis (22), including a subset of the two large cohorts of current and former smokers from the SPIROMICS and COPDGene cohorts, protein-Quantitative Trait Locus (pQTL) approaches were used to identify single nucleotide polymorphisms (SNPs) associated with measurement of 88 blood proteins, including E-cadherin. Analysis of both SPIROMICS and COPDGene suggests a locus on chromosome 19 as a key determinant of serum E-cadherin regulation (**Fig 1A-B**), which our lab found strongly correlates with lung epithelial E-cadherin in a group of patients at risk for COPD (**Supplementary Figure 1**). The peak SNP rs516246 (meta-analysis, p=4x10^-27^) is the strongest locus in both cohorts with a p-value of 8.95x10^-16^ and 1.21 x10^-16^ in SPIROMICS and COPDGene, respectively. This is a trans pQTL for E-cadherin as the locus maps to an intronic region in *FUT2,* the gene encoding fucosyltransferase-2 (FUT2). Since we did not have FUT2 protein levels in these patients, we leveraged results from the GTEx consortium (23) to look for evidence of transcript regulation of FUT2 levels by these SNPs. We found that rs516246 is a significant eQTL for *FUT2* transcript levels (**Fig 1C**) across numerous tissues, but notably, also in lung. Individuals heterozygous for this eQTL have decreased levels of *FUT2* mRNA transcript. When homozygous, transcript levels are found to be even less. This revealed the peak pQTL SNP is in 100% linkage disequilibrium with an exonic SNP rs601338, previously shown to result in no expression of the fucosyltransferase-2 (24).

**Figure 1:**
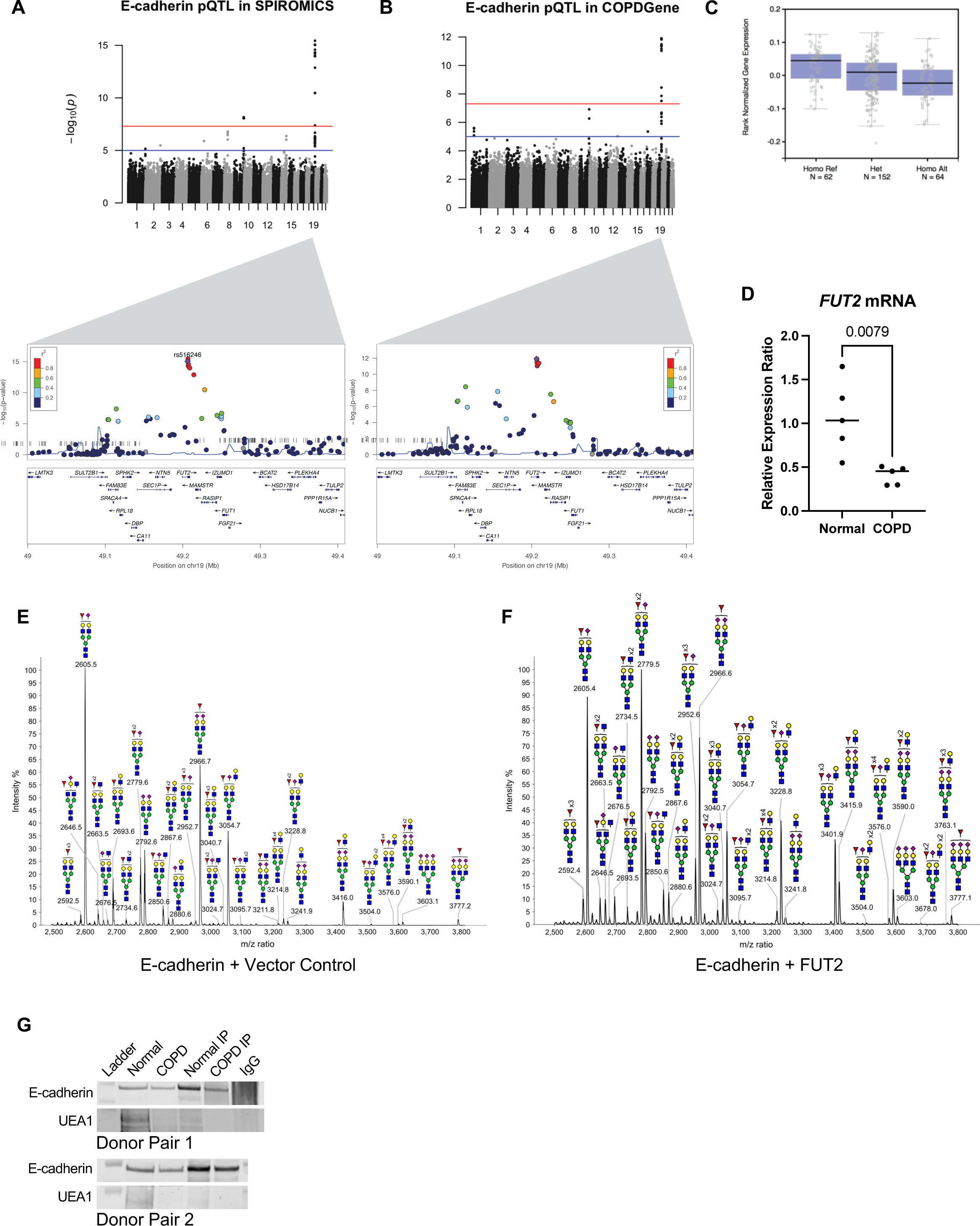
Fucosyltransferase-2 fucosylated E-cadherin. **A.** Protein quantitative trait loci (pQTL) analysis of E-cadherin variants in the SPIROMICS database. **B.** Protein quantitative trait loci (pQTL) analysis of E-cadherin variants in the COPDGene database. **C.** Measured expression quantitative trait loci (eQTL) transcript levels of FUT-2 by rs516246 genotype in lung tissue from Genotype-Tissue Expression (GTEx). **D.** *FUT2* mRNA transcript is significantly decreased in COPD derived bronchial epithelia (*p = 0.0079*). **E.** Partial MALDI-TOF mass spectra (m/z 2500-3800) of E-cadherin N-glycans. **F.** Partial MALDI-TOF mass spectra (m/z 2500-3800) of E-cadherin co-expressed with FUT2 N-glycans, demonstrating an increase in fucosylation. **G.** An immunoprecipitation of E-cadherin indicates less UEA-1+ E-cadherin in bronchial epithelial cells derived from COPD patients.

### E-cadherin is fucosylated by fucosyltransferase-2

Given the strong genetic association between *FUT2* and *CDH1* in two independent cohorts, we found it pertinent to determine whether we could establish a functional relationship. Based on previous literature there are four N-glycosylation sites in E-Cadherin at residues 554, 566, 618, and 633 (25). We utilized MALDI-TOF mass spectrometry based glycomic methodologies to characterize the N-glycans of E-Cadherin when co-expressed with FUT2. A heterogeneous glycan profile is observed which is dominated by complex type N-glycans with fucosylation and/or sialylation (m/z 1590-4588, NeuAc_0-4_Gal_0-4_Man_3_GlcNAc_4-6_Fuc_0-5_). More minor levels of high mannose glycans (m/z 1579-2396, Man5-9GlcNAc2) are also observed (**Supplementary Figure 2**) by comparing the relative intensities of related fucosylated N-glycan molecular ions levels of E-Cadherin fucosylation increase when co-expressed with FUT2. For example, signals at m/z 2605, 2779 and 2952 which are consistent with a monosialylated bi-antennary complex glycan with 1, 2 and 3 fucose residues and signals at m/z 3054, 3228, 3401 and 3576 which are consistent with a monosialylated tri-antennary complex glycan with 1, 2, 3 and 4 fucose (Fuc) residues (**Fig 1D-E**).

**Figure 2:**
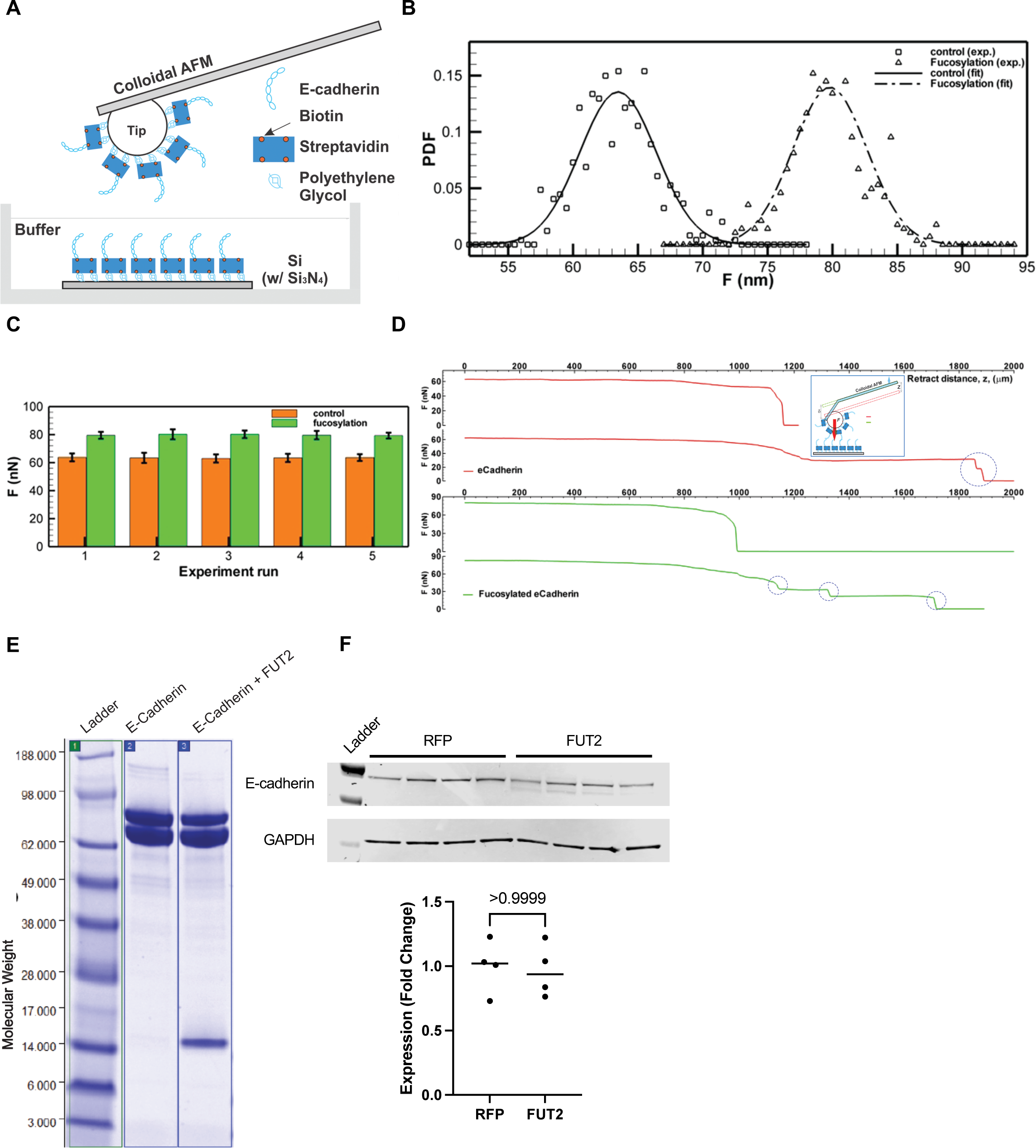
FUT2-fucosylated E-cadherin has higher bond strength than non-FUT2-fucosylated E-cadherin. **A.** Schematic of the experimental setup for AFM based nano-indentation measurements of e-cadherin with or without fucosylation. **B.** Probability Density Function (PDF) of maximum adhesion forces between e-cadherin filaments. Square: non-fucosylated e-cadherin. Delta: fucosylated e-cadherin. Lines: normal distribution fits for non-fucosylated (solid: *F̄* = 63.47 *nN, σ* = 2.8585 *nN*) and fucosylated e-cadherin (dashed: *F̄* = 79.801 *nN, σ* = 2.9432 *nN*) **C.** Bar graph showing mean maximum adhesion force (nN) of non-fucosylated (dark) and fucosylated (light) e-cadherin. Error bar: one standard deviation. Total five pairs of experimental runs are presented. (Each run contains ∼100 measurements, **Supplementary Table 1**) **D.** Sample adhesion curve showing the maximum binding force and break-offs (cliffs, blue circle). Single (Row 1) and two (Row 2) break-offs of the bonded non-fucosylated e-cadherin filaments, while Row 3 & 4 show maximum & partial break-offs of the bonded fucosylated e-cadherin filaments. Inset: schematics of adhesion measurement using AFM. **E.** Immunoblot depicting the presence of N-terminal fragment of E-cadherin when co-expressed with *FUT2*. **F.** E-cadherin expression does not change with FUT2 overexpression.

More detailed N-glycan structural analysis, in particular the assignment of the positions of fucosylation, was achieved by MS/MS analysis of selected molecular ions. Exemplar data is shown for the m/z 2779 molecular ion with composition of NeuAc_1_Gal_2_Man_3_GlcNAc_4_Fuc_2_. Key fragment ions which indicate the fucosylation of the terminal Gal residue, and therefore indicate the action of FUT2, include m/z 433, 834 and 1967. All these ions increased abundance when E-Cadherin was co-expressed with FUT2 (**Supplementary Figure 2**).

### Fucosylated E-cadherin has higher E-cadherin-E-cadherin bond strength

Terminal fucosylation of E-cadherin by FUT2 occurs in the extracellular domain, the region of the protein that mediated bonds between E-cadherin molecules, and increased E-cadherin bonds increase surface stabilization of the protein. Therefore, we sought to determine if FUT2-dependent fucosylation affected bond strength. We measured the protein bond strength of purified cell-free FUT2-fucosylated E-cadherin or E-cadherin control using atomic force microscopy (AFM). (**Fig 2A**). We found that FUT2-fucosylation increased the bond strength of E-cadherin. Plotting the probability density functions of force measurements revealed that FUT2-dependent fucosylation resulted in distinct force distributions (**Fig 2B**). FUT2-dependent fucosylation increased E-cadherin bond strength from ∼63 nN (without terminal fucosylation) to ∼80 nN (with terminal fucosylation, **Fig 2C**). Of note, although the strength was higher with FUT2-dependent fucosylation, we were able to detect more break-off events of E-cadherin when compared to unfucosylated-E-cadherin (**Fig 2D**). A Coomassie stain confirmed purified protein for each condition (**Fig 2E**). It is interesting that in the fucosylated E-cadherin, an additional N-terminal fragment which was verified by mass spectrometry to be an extracellular fragment of E-cadherin isolated along with the purified protein, suggesting the high adhesion of the extracellular domain. It is of note that FUT2 did not change the abundance of E-cadherin (**Fig 2F**) or CDH1 transcript (**Supplementary Figure 3**) but did result in the presence of an additional band with a slightly lower molecular weight in A549s.

**Figure 3:**
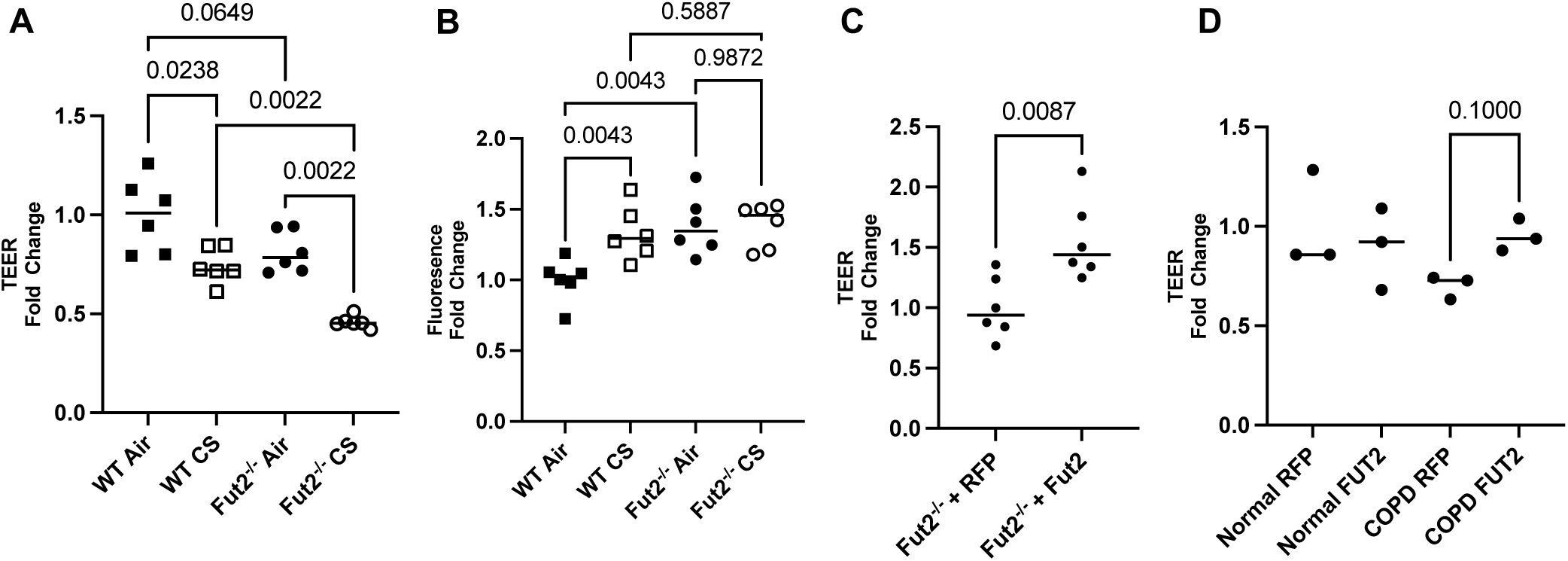
FUT2 is necessary to maintain integrity of the airway epithelium through its regulation of E-cadherin. **A.** *Fut2^-/-^* mTEC have lower TEER than WT mTEC, which is worsened by CS-induced injury (n = 6). **B.** *Fut2^-/-^*mTEC have higher permeability than WT mTEC, but it is not further worsened by CS exposure (n = 6). **C.** Reconstitution of *Fut2* in *Fut2^-/-^*mTEC recovers TEER (n = 6). **D.** Overexpression of FUT2 improves TEER in COPD derived epithelia (n = 3).

### Fut2 is required to maintain epithelial barrier function

At baseline exposure to air, Fut2 deficient mouse tracheal cells (mTEC) show decreased epithelial integrity as measured by decreased transepithelial electrical resistance (TEER) and increased FITC dextran flux (**Fig 3A-B**). CS induced a decrease in TEER in both WT and *Fut2^-/-^* mTEC. Interestingly, while CS induced a significantly increased permeability in WT mTEC, this worsening barrier was not noted in *Fut2^-/-^* mTEC suggesting that a loss of Fut2 is sufficient alone to cause epithelial barrier dysfunction. Lentiviral mediated restoration of Fut2 shows increased epithelial integrity, evident by increased TEER compared to empty vector controls (RFP) and nearing that of WT cells (**Fig 3C**). As a proof of principle, we demonstrate that overexpression of FUT2 also increases TEER in COPD derived bronchial epithelial cells (**Fig 3D**). Transduction efficiency is demonstrated in **Supplementary** Figure 4.

### Lack of Fut2 results in increased susceptibility to elastase induced emphysema and fibrosis

Elastase is commonly used as an *in vivo* model for emphysema (15, 26, 27). Representative 10X and 0.5X H&E images reveal that after intratracheal administration of elastase in WT and *Fut2^-/-^* mice there was marked alveolar destruction. However, *Fut2^-/-^* mice developed visibly worse emphysema (**Fig 4A**). Masson’s Trichrome staining demonstrates increased collagen deposition in *Fut2^-/-^* airways and alveoli, both more apparent in mice treated with elastase (**Fig 4B**). *Fut2^-/-^* mice have increased total lung capacity and residual volume compared to the WT mice (**Fig 4C-D**). Compliance increased with elastase treatment in WT mice but demonstrated no significant change in *Fut2^-/-^* mice (**Fig 4E**). However, this is consistent with our finding of both *Fut2* airspace enlargement and increased fibrosis on lung histology (**Fig. 4B**). Elastase induced a significant increase in mean linear intercept in *Fut2^-/-^*, but not WT mice (**Fig 4F**). *Fut2^-/-^* was confirmed by qPCR (**Fig 4G**).

**Figure 4:**
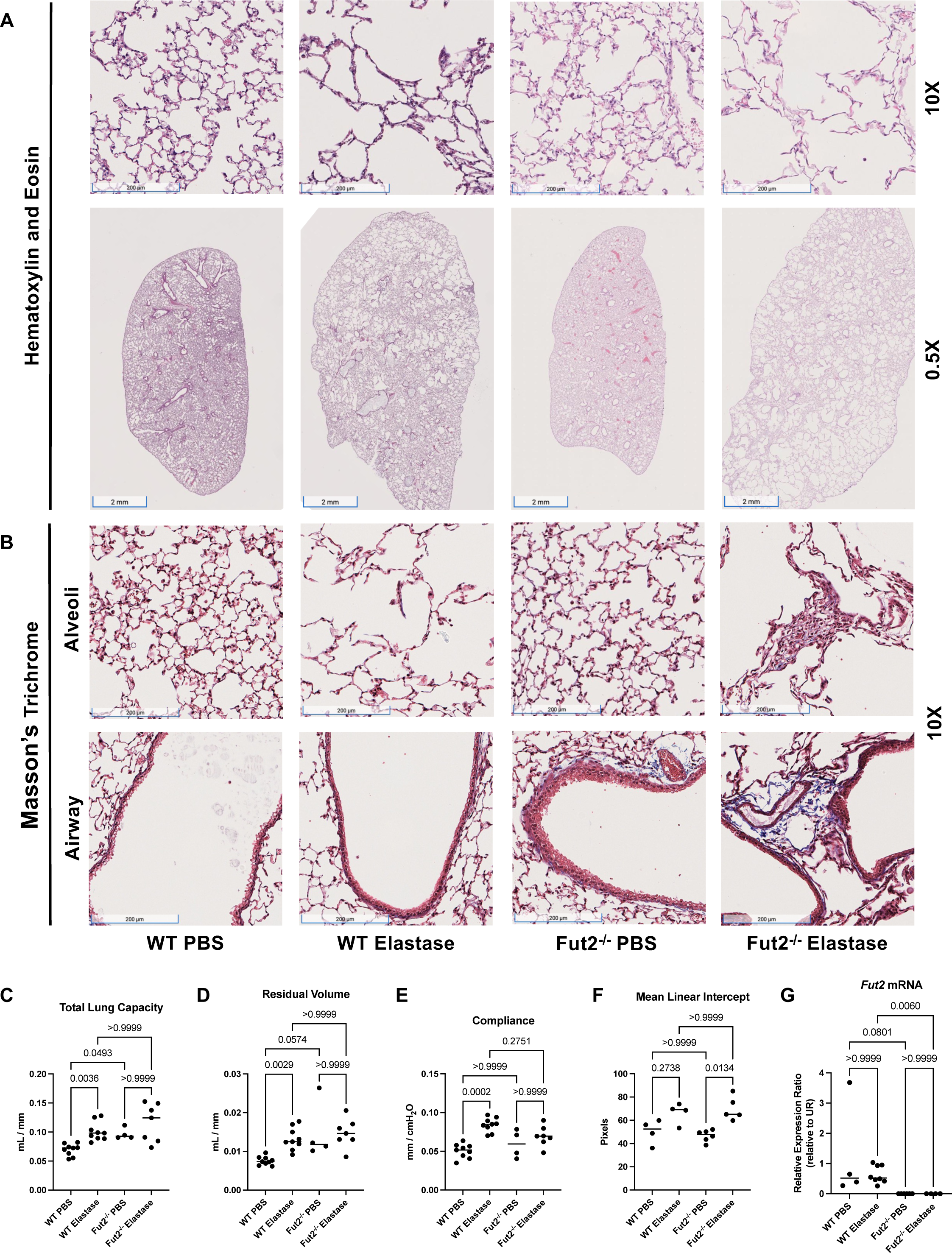
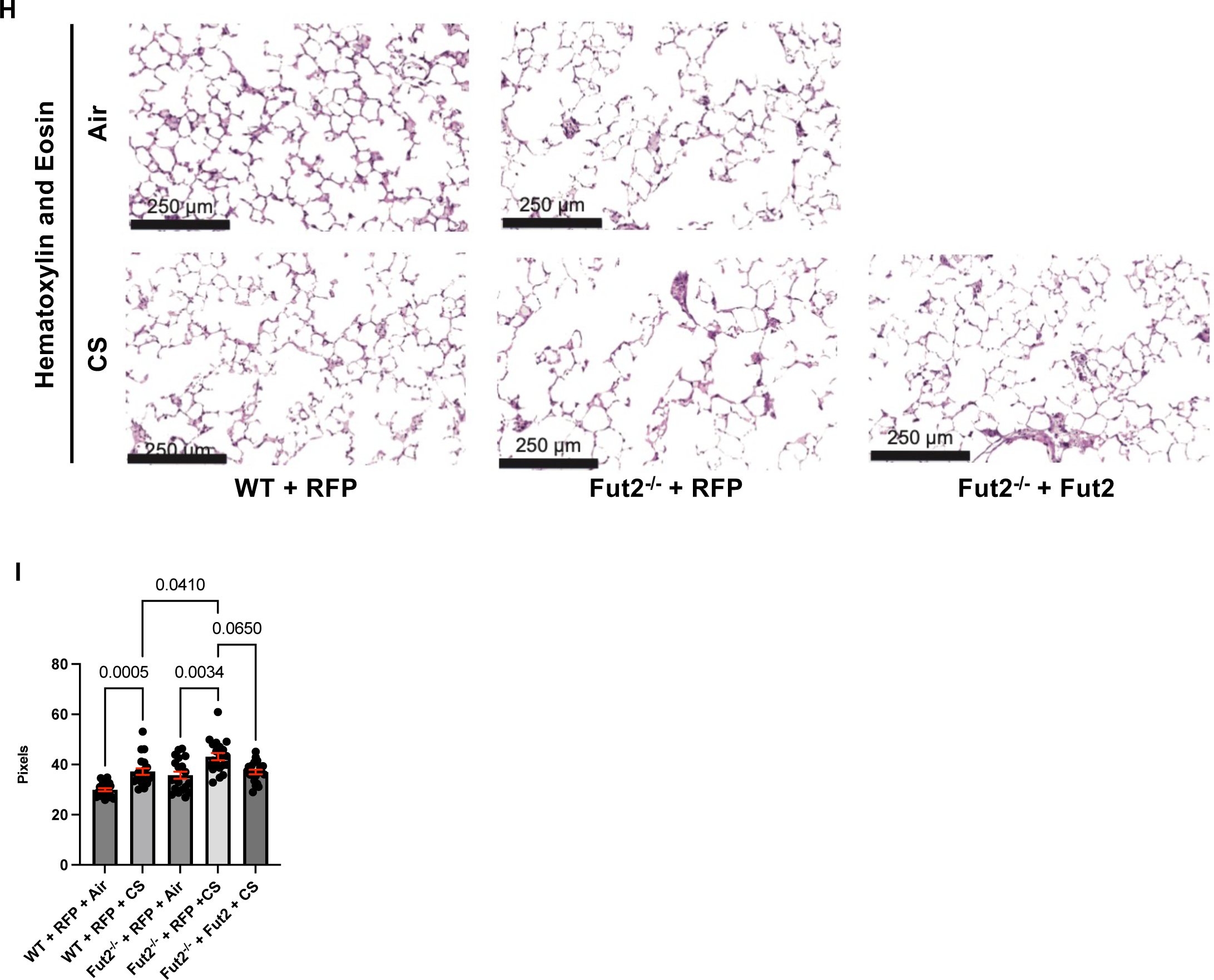
*Fut2^-/-^*mice develop fibrosis and emphysema with elastase administration. **A.** Hematoxylin and eosin staining of WT and *Fut2^-/-^* mice administered PBS or elastase intratracheally (n = 3). **B.** Masson’s Trichrome staining indicates increased fibrosis of the alveoli and airways in *Fut2^-/-^*mice (n = 3). **C.** *Fut2^-/-^* mice have increased total lung capacity and **D.** residual volume (n = 4 to 9). **E.** Compliance does not significantly change in *Fut2^-/-^* mice (n = 4 to 9). **F.** Elastase induces alveolar destruction, measured by mean linear intercept (n = 4 to 6). **G.** *Fut2^-/-^* mice have undetectable *Fut2* transcript in the lung (n = 4-8). **H.** Hematoxylin and eosin staining of of WT and *Fut2^-/-^*PCLS exposed to air or CS. **I.** *Fut2^-/-^* PCLS show increased susceptibility to CS as measured by mean linear intercept, which is rescued by reconstitution of *Fut2*.

### Fut2 deficiency increases susceptibility of precision cut lung slices to CS

Using mouse precision-cut lung slices, we have demonstrated *Fut2* expression modifies CS-induced alveolar destruction. CS exposure of WT PCLS transduced with a control vector (RFP) increased the mean linear intercept (MLI) indicating alveolar destruction. At baseline the *Fut2-/-* PCLS demonstrated an increased mean linear intercept compared to the background WT mice. Moreover, *Fut2-/-* RFP PCLS exposed to CS showed a significantly higher MLI than WT RFP PCLS exposed to CS. However, when *Fut2* is reconstituted in *Fut2-/-* PCLS, it abrogates CS induced injury (**Fig 4I-J**). Interestingly, unlike the mouse elastase model, the PCLS did not demonstrate increased fibrosis, which could be a limitation of the model. Transduction efficiency is demonstrated in **Supplementary Figure 4.**

### FUT2 is necessary for sufficient proliferation of the epithelium

E-cadherin has mostly been studied in the context of contact inhibition, although there is some evidence that E-cadherin can promote cell proliferation in some cancer models (7, 15, 28). Previously we have shown that a loss of E-cadherin results in decreased proliferation in lung basal cells (15). Overexpression of FUT2, in the lung epithelial cell line A549, results in increased proliferation as measured by doubling time and KI67+ cells (**Fig 5A-C**). The proliferation defect resulting from knockdown of E-cadherin, a phenocopy of COPD, is recovered by overexpression of FUT2 (**Fig 5D-E**).

**Figure 5:**
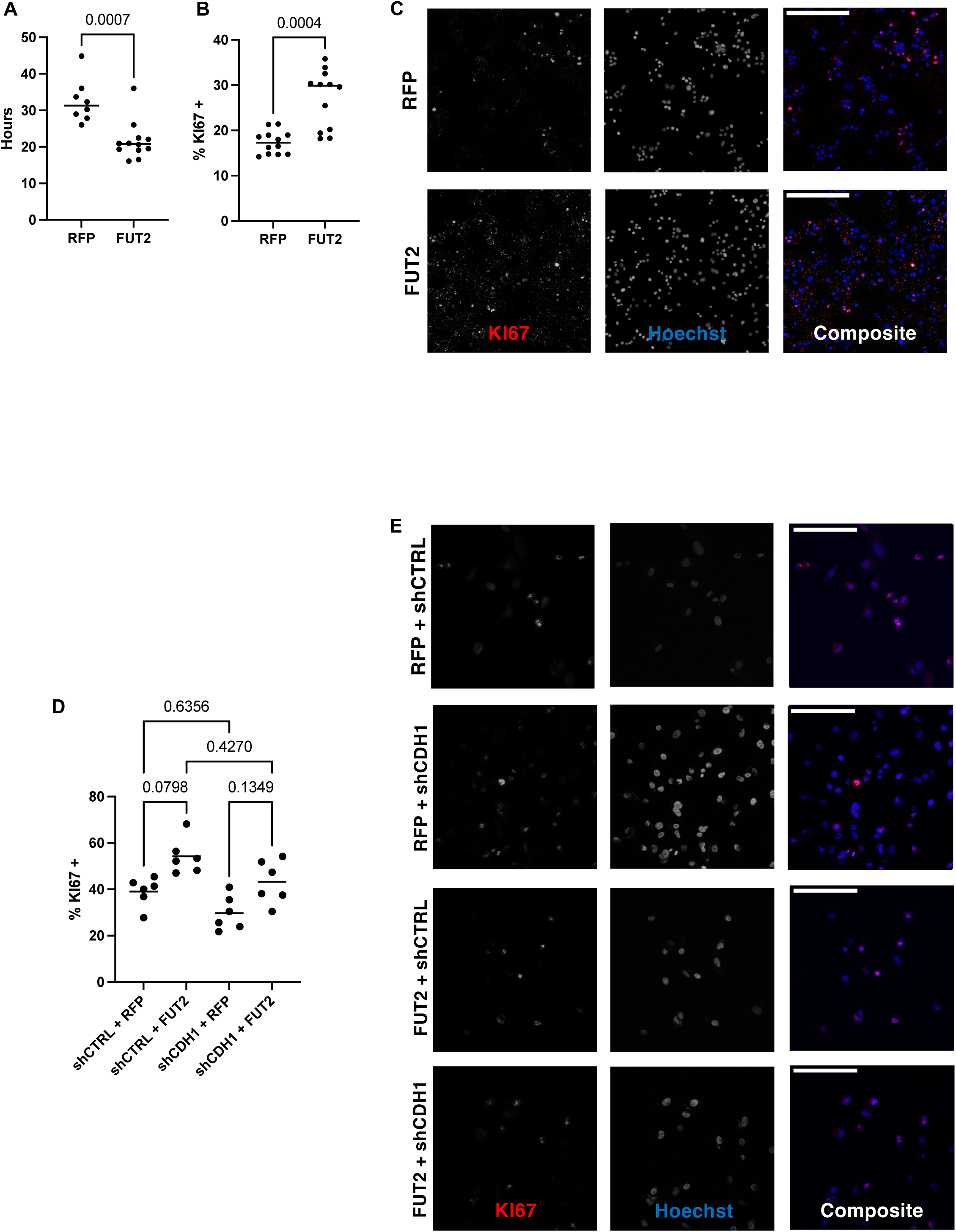
FUT2 is required for sufficient proliferation of the lung epithelium. **A.** Overexpression of FUT2 decreases doubling time of A549s (n = 8 to 12). **B and C.** Overexpression of FUT2 increases the number of KI67+ cells (n = 12); scale bar (upper left) = 250 microns. **D and E.** Knockdown of CDH1 decreases the number of KI67+ cells, overexpression of FUT2 in CDH1 knockdown A549s has a limited effect (n = 6); scale bar (upper left) = 125 microns.

### FUT2 is required for a well-differentiated airway epithelium

Given E-cadherin’s central role in the maintenance of a well differentiated epithelium, we examined cell type specific markers for club cells (*Scgb1a1*), goblet cells (*Muc5ac*), basal cells (*Krt5*), and ciliated cells (*Foxj1*). This revealed both the club cell and ciliated cell populations are dependent on *Fut2* expression *in vivo* (**Fig 6A-D**). Interestingly, this did not correlate with our human *in vitro* data (**Fig 6E-H**).

**Figure 6:**
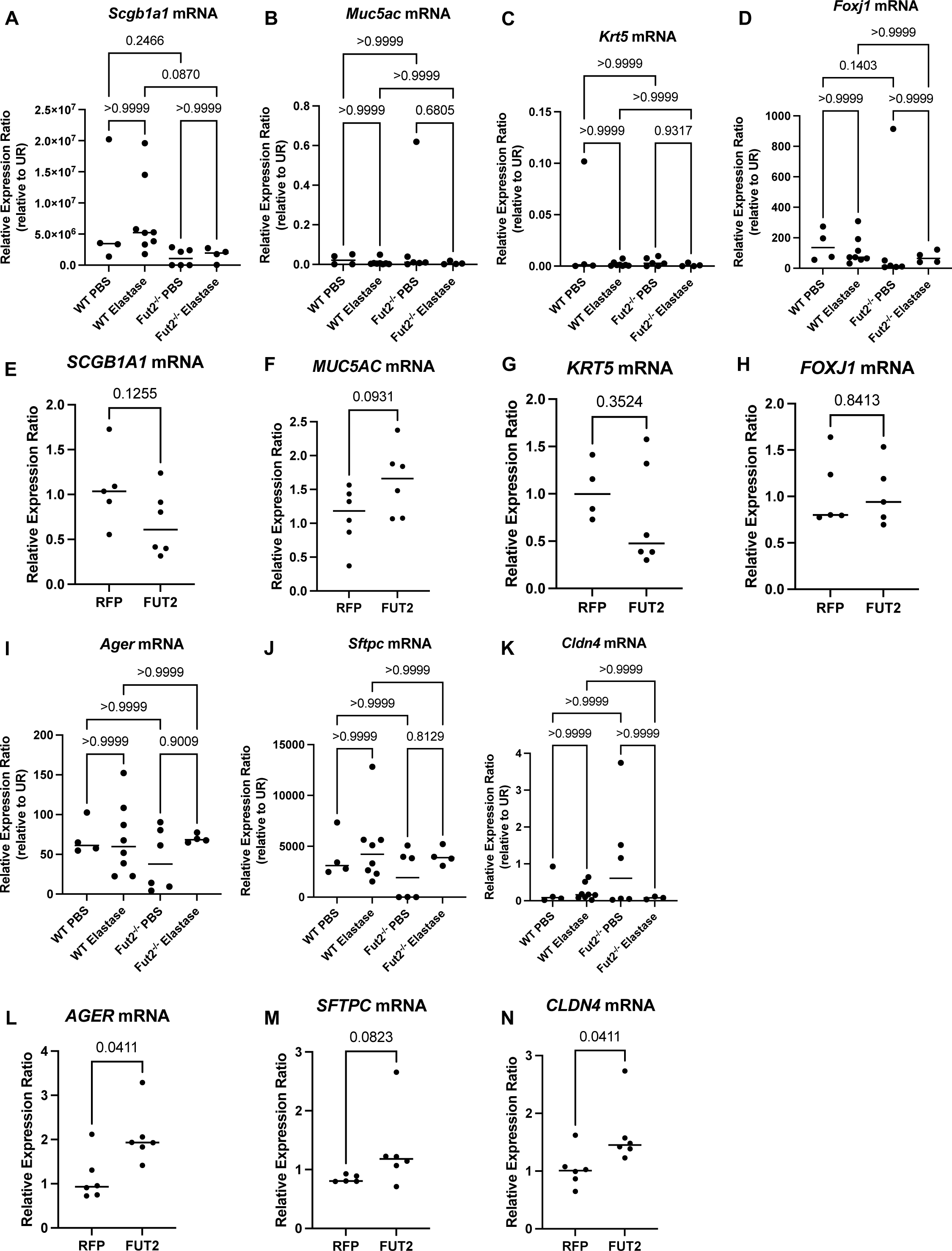
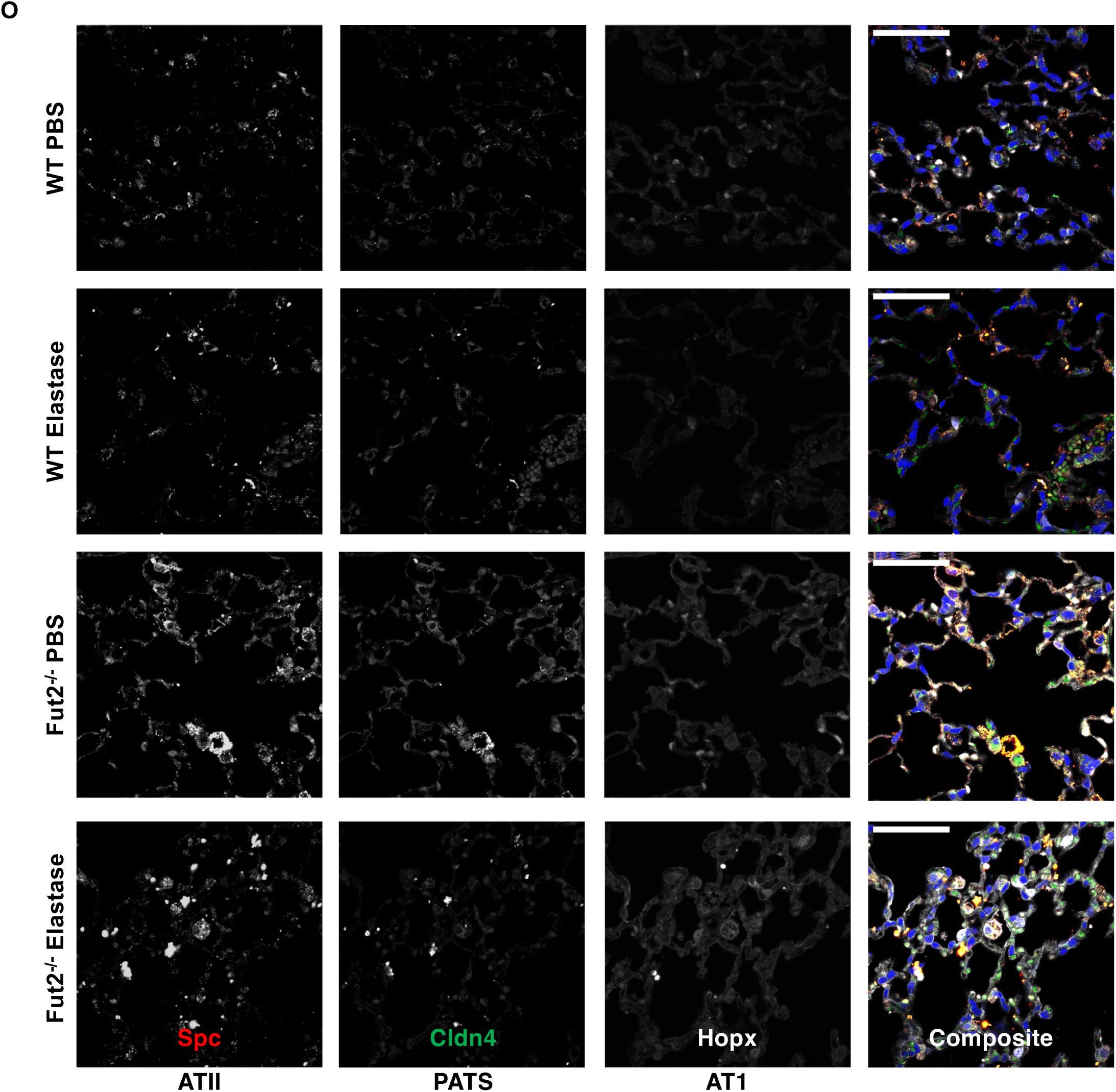
FUT2 is required for regeneration of the alveolar and airway epithelia. **A.** *Scgb1a1* transcripts are decreased in *Fut2^-/-^* mice instilled with elastase when compared to WT. **B and C.** *Muc5ac* and *Krt5* transcripts are unchanged with loss of *Fut2* or elastase. **D.** *Foxj1* expression trends to be lower with loss of *Fut2*. **E.** *SCGB1A1* expression slightly decreases with FUT2 overexpression. **F.** *MUC5AC* expression slightly increases with FUT2 overexpression. **G and H.** *KRT5 and FOXJ1* expression are relatively unchanged with FUT2 overexpression. **I, J, and K.** There is no change in *Ager, Sfptc,* or *Cldn4* expression with loss of *Fut2* or elastase. **L, M, and N.** *AGER, SFTPC, and CLDN4* expression increase with FUT2 overexpression. **O.** *Fut2^-/-^* mice have increased Spc+ cells in both PBS and elastase. Elastase induces increased Cldn4 expression (n = 2); scale bar (upper left) = 50 microns.

### FUT2 is required for ATII to AT1 transdifferentiation

Analysis of RNA from whole lung homogenate from WT and *Fut2^-/-^* mice instilled with PBS or elastase indicates no significant change in *Ager* (AT1)*, Sftpc* (Spc, AT2), or *Cldn4* (PATS) expression (**Fig 6I-K**). However, overexpression of *FUT2* resulted in increased the transcriptional expression of *AGER, SFTPC,* and *CLDN4* compared to the empty vector control (RFP) (**Fig 6L-N**) suggesting increased activation of the AT2 to AT1 transdifferentiation pathway. Analysis of protein via immunofluorescence revealed a stark difference in the abundance of various cell types (**Fig 6O**). In *Fut2^-/-^* mice there is a distinct increase in Spc+ cells indicating a persistence of alveolar type II (AT2) cells when compared to WT. Elastase resulted in increased Cldn4 expression (PATS cells) in both WT and *Fut2^-/-^*mice, but there is notably more present in *Fut2^-/-^* mice. Hopx+ (AT1) cells remained nearly constant in WT vs *Fut2^-/-^* mice in both PBS and elastase conditions.

## Discussion

Genome-wide association studies have associated lower *CDH1* levels in COPD patients with a worsened prognosis (9, 29–31) and our lab has demonstrated that genetic loss of E-cadherin alone is sufficient to cause both airspace enlargement and airway disease (15). However, to date there has not a body of literature exploring the molecular mechanisms by which E-cadherin is regulated in the context of COPD. This is difficult due to the large clinical variability, however dissecting mechanisms mediating increased susceptibility and identifying strategies to reduce it accordingly can provide the necessary base of knowledge for both risk stratification and therapeutic intervention. While the obvious answer may be to increase E-cadherin abundance, this can be technically challenging and regulation of protein abundance is a very small aspect of maintenance of the proteome. We demonstrate that improving function of E-cadherin is also sufficient providing another potential therapeutic avenue for a multipronged approach targeting COPD.

We have identified a novel post-translational modification of E-cadherin namely terminal fucosylation catalyzed by fucosyltransferase 2. This significantly increases bond strength between E-cadherin molecules, which is critical given E-cadherin bond strength is correlated with protein stability (32, 33). Due to experimental limitations, it is unclear whether this increased bond strength is due to increased aggregation of E-cadherin or because single molecules of E-cadherin have strong trans-bond strength. As the AFM probe, a SiO_2_ sphere, allows for multiple E-cadherin molecules to attach meaning both possibilities would result in increased observed bond strength. Although future studies with a smaller probe could differentiate between these possibilities, both mechanisms result in the desired result of increased adhesion. In addition to increased bond strength, it is interesting that the fucosylated E-cadherin had several breaks before the full bonds separated. As there were several identified fucosylation sites, with the potential of differential fucosylation of E-cadherin with the overexpression of FUT2, it is possible that these breaks reflect the disruption of E-cadherin with fewer fucosylated sites. Future studies may be able to dissect this possibility further.

Our data, both *in vitro* and *in vivo*, clearly demonstrate a significant impact on regeneration and repair of the airway and alveolar epithelia. We show that epithelial barrier integrity requires FUT2 and propose that defects in proliferation and differentiation are the cause. We found there is a clear dependence of proliferation on FUT2 expression, with proliferation increasing with FUT2 expression. In addition to having a significant effect on proliferation, we also demonstrate FUT2 is required for a well-differentiated airway and alveolar epithelium. Of note, loss of Fut2 results in accumulation of Spc+ (AT2) cells, which is particularly interesting considering we have previously shown knockout of Cdh1 in Spc+ cells alone is sufficient to cause emphysema (15). This begs the question of the fate of Spc+ cells that lack Cdh1 and whether they can effectively transdifferentiate into alveolar type 1 cells.

Using large clinical datasets, we demonstrate a significant genetic association between *CDH1* and *FUT2* prevalent in smokers, former smokers, and COPD patients, confirming previous literature (22). Here we extend this genetic association with an arsenal of molecular approaches, ultimately determining this genetic association extends to a molecular interaction with far reaching impacts holding significant therapeutic potential.

## Methods

### Animals and study design

This study was approved by the Institutional Animal Care and Use Committee of the Johns Hopkins University Animal Use and Care Committee and compiled within the Guidelines for Care and Use of Laboratory Animals issued by the National Institutes of Health. The study used both C57BL/6 (Jackson Laboratory, Bar Harbor ME) and *Fut2^-/-^* (kindly donated by Dr. Christopher Evans, University of Colorado Anschutz) strains. All mice were bred and maintained in a specific pathogen-free environment.

### pQTL and eQTL Results

To identify pQTLs for E-Cadherin protein levels, we leveraged previously published QTL genomewide scans from two large cohorts of current and former smokers with and without COPD [SPIROMICS (N = 750); COPDGene (N = 590)]. As previously described, the study aimed to identify single nucleotide polymorphisms (SNPs) associated with measurement of 88 blood proteins (protein quantitative trait loci; pQTLs) including E-Cadherin. Here, we extracted the genomewide QTL analysis for E-Cadherin alone. To identify what gene expression the identified pQTLs for E-Cadherin were associated with, we leveraged the GTEx portal (https://www.gtexportal.org/home/).

### FUT2 treatment on E-cadherin

E-cadherin with 6x His-tag and native FUT2 enzyme synthetic genes were created. After plasmid DNA was purified, E-cadherin DNA was transfected either with or without FUT2 DNA in EXPI293 cells. E-cadherin was then purified from both conditions with nickel capture and size exclusion chromatography.

### Mass Spectrometry

N-linked glycan analysis was performed according to Jang-Lee et al (34). The E-Cadherin glycoprotein samples were reduced, carboxymethylated, and digested with trypsin. N-Glycans were enzymatically released by peptide N-glycosidase F (E.C. 3.5.1.52; Roche Applied Science) digestion then purified by C18-Sep-Pak (Waters Corp., Hertfordshire, UK). The purified *N*-glycans were permethylated using the sodium hydroxide procedure and purified by C18-Sep-Pak. The permethylated *N*-glycans were then dissolved in methanol before an aliquot was mixed at a 1:1 ratio (v/v) with 10 mg/ml 3,4-diaminobenzophenone in 75% acetonitrile. The glycan-matrix mixture was spotted on a stainless-steel target plate and dried in vacuum. MALDI-TOF MS and MALDI-TOF/TOF MS/MS data were obtained using a 4800 MALDI-TOF/TOF mass spectrometer (AB Sciex UK Limited) in the positive-ion mode. For MS/MS, the collision energy was set at 1 kV, and argon was used as the collision gas. The obtained MS and MS/MS data were viewed and processed using Data Explorer 4.9 (AB Sciex UK Ltd). All N-glycans were assumed to have a core of Manα1-6(Manα1–3)Manβ1–4GlcNAcβ1–4GlcNAc based on known biosynthetic pathways and susceptibility to peptide N-glycosidase F digestion. Monosaccharide compositions in terms of numbers of Hex, HexNAc, etc. derived from MALDI-MS. MALDI-TOF/TOF MS/MS fragment ions were identified manually and with the assistance of the Glycoworkbench tool (35).

### Immunoprecipitation (IP)

1000 ug of protein, determined by BCA assay were precleared for 1 hour with Protein G Sepharose 4 Fast Flow beads (GE17-0168-01, Millipore Sigma) at 4°C. Following preclearance, protein lysate was removed and coupled with a polyclonal E-Cadherin antibody (20874-1-AP, Proteintech Group, IL, USA) overnight at 4°C. Antibody coupled lysate was then incubated with Protein G Sepharose 4 Fast Flow beads for 1 hour at 4°C. Supernatant was removed for further analysis and RIPA, 4X Bolt LDS Sample Buffer (B0007, ThermoFisher Scientific), and 10X Bolt Sample Reducing Agent (B0009, ThermoFisher Scientific) were added to the Sepharose beads. Samples were then boiled at 95°C for 15 minutes with agitation. The supernatant, which contained the immunoprecipitated protein, was then removed to be analyzed via western blot.

### Western blot assay

Western blot analysis was carried out as previously described (15, 36). Briefly, proteins were separated on Bolt 4 – 12%, Bis-Tris gradient gel (ThermoFisher Scientific, NY, USA) and then transferred to a Immobilion-P PVDF membrane (Millipore Sigma, MA, USA). Following transfer, the PVDF membrane was blocked in 5% w/v BSA in 1X PBS with 0.1% Tween® 20 Detergent, (1X PBST, Millipore Sigma, MA, USA). The membrane was then probed for E-cadherin, GAPDH, and UEA1 (E-cadherin (24E10) Rabbit mAb, 135 kDa and GAPDH (14C10) Rabbit mAb, 37 kDa antibodies from Cell Signaling Technology, MA, USA), Ulex Europaeus Agglutinin I Biotinylated (B-1065-2) from Vector Laboratories) according to manufacturers’ instructions. The blot was then probed with secondary antibody (IRDye® 800CW Streptavidin and IRDye® 680RD Goat anti-Rabbit IgG Secondary Antibody, LI-COR) and imaged. Blots were quantified using ImageStudio (LI-COR).

### Quantitative Polymerase Chain Reaction (qPCR)

RNA was extracted from human bronchial epithelial cells / mice lung tissues and purified Trizol (ThermoFisher). RNA was then converted to cDNA after addition of dNTP mix, 10X RT Random Primers, Reverse Transcriptase, and nuclease free water and the following PCR cycle: 25°C for 10 minutes, 37°C for 120 minutes, 85°C for 5 minutes (High Capacity cDNA Reverse Transcription Kit, ThermoFisher). Following conversion, equal amounts of cDNA from each sample were added to previously designed primers and SYBR Green mix in duplicate. The following qPCR cycle was run: 95°C for 10 minutes, 40 cycles of 95°C for 15 seconds, 60°C for 1 minute. Relative expression ratio was determined using the method. Based on comparative Ct method, gene expression levels were calculated utilizing GAPDH as the housekeeping gene. Primers are listed in **Supplementary Table 2.**

### Atomic Force Microscopy (AFM)

E-cadherin was conjugated directly over a substrate of 15*mm* × 15*mm* n-type silicon wafer fragment and a gold coated AFM colloidal probe (HQ:CSC38/Cr-Au, MikroMasch). A 4” silicon wafer was coated with silicon nitride (*si*_3_*n*_4_) by a PECVD (Plasmatherm 790) at 250 °C for 5min to achieve a 100nm thick *si*_3_*n*_4_ thin film. The wafer was then diced into a series of 15*mm* × 15*mm* fragments for experimentation.

### Probe and surface preparation

Before functionalization, both substrate and probe were cleaned with piranha etching solution (*H*_2_*O*_2_:*H*_2_*SO*_4_ at 1:2 w/w). While a 15*mm* × 15*mm* substrate was cleaned in freshly prepared solution for 30min, the gold coated AFM probes, held in an in-house made HDPE probe holder, were cleaned with one-day old cold piranha solution for 10 seconds. Note that probes remained in the holder for cleaning and later functionalization procedure. After cleaning, the testing substrate was subsequently rinsed with DI water, acetone, methanol, isopropanol, and then DI water again, while probes were soaked in abovementioned solutions for >10 minutes each with gentle shaking. The testing substrate was then dried with ’_#_and baked at 120 ℃ for 1 min. The AFM probe was air dried in a glove box (EW-34788-10, Cole-Parmer) with N_2_ purging overnight.

### Probe and surface functionalization

Illustrated with **Fig. 4A**, to attach E-cadherin to the surfaces of substrates and probe, we applied amino functionalization by APTES (3-aminopropyltriethoxysilane) to the probe and wafer. Before amino functionalization, the stock APTES (CAS 919-30-2, Sigma-Aldrich) was purified at the distillation temperate of 103°C under a vacuum of 20mmHg. The purified APTES was then dispensed into 1-2ml screw cap vials in an Ag filled glove box. Shelf-life for the distilled APTES was over 6 months when stored at -20 °C.

### Aminosilanization

Probe and wafer were functionalized by a gas phase aminosilanization procedure. 30 *μl* of APTES and 10 *μl* of triethylamine were placed in two separate trays together with probe and testing wafer into a desiccator. The desiccator was pumped down to 200 mTorr and then filled to 75 Torr with Argon. Vapor deposition continued for 4 hours. The APTES on probe and wafer was cured at room temperature in an Argon filled glove box for 2 days.

### APTES conjugation

Probe and wafer surfaces was further functionalized by a biotinylated polyethylene glycol (PEG). PEGylation solution of 80mg of m-PEG-SVA (5 kDa, M-SVA-5K, Laysan Bio, Inc) and 4 mg of Biotin-PEG-SVA (Laysan Bio, Inc.) in 320,-PEGylation buffer (0.1M sodium bicarbonate) was freshly prepared. A make-shift reaction chamber made from a 5” petri dish was used to perform PEGylation functionalized as following: A small tray containing 2 ml of DI water was placed in the chamber to maintain the humidity. The wafer fragment was placed in the reaction chamber with the functionalized surface facing up. A 70,-of the PEGylation mixture was deposited over each testing wafer fragment (e.g. 15 mm×15mm). To prevent the cantilever from being destroyed, the probe functionalization was completed by dipping the probe in the mixture mounted on an in-house developed mini-manipulator. The apparatus including probe, manipulator, and PEGylation container was sufficiently small to be enclosed in a 5” petri dish. The chamber was then sealed with parafilm and placed in dark overnight. After functionalization, the testing wafer fragments were then rinsed with DI water and dried with N_2_; while the probe mounted on manipulator was soaked in DI water bath for 20 minutes and then air dried in N2 filled glovebox. The PEGylated cantilever and testing wafer surfaces can be stored in desiccator for ∼2 weeks. Note that most of abovementioned procedures were performed in a laminar hood.

### Streptavidin attachment

The PEG-functionalized testing surface and probe were incubated in 0.1 mg/ml BSA in TB buffer (i.e. 10 mM Tris, 100 mM NaCl, 10 mM KCl, and 2.5 mM CaCl_2_) for 12 hours to minimize nonspecific protein binding, followed by the incubation with 0.1 mg/mL streptavidin (Sigma-Aldrich) in TB buffer for 30 min.

### E-Cadherin immobilization

The surface and probe were incubated in 200 nM biotinylated E-cadherin (Sinobiological, Inc) in TB buffer for 45 min. After E-cadherin immobilization, the surface and probe was further incubated in 2 µM biotin in TB buffer for 10 min and the free biotin was washed away using TB buffer.

### AFM measurement

After functionalization, the testing surface and probe were mounted in environmental AFM (AFMWorkshop, LLC). The testing surface was placed in a flow cell containing TB buffer. Each wet nano-indentation measurement was performed by a procedure. To promote the interaction of E-cadherins between surface and probe, the probe was allowed to extend 200nm into the surface at the rate of 500nm/s. At the end of the extension, the probe and surface maintained in contact for 10min with nominal contact force of ∼5nN. After 10min “binding” period, the probe was allowed to retract 13,# away from the surface at the rate of 500nm/s. The large retraction distance was specifically selected to ensure the separation of the probe and surface. The above procedure was repeated at different locations (e.g. totaling ∼100 per experiments). Note that these locations were randomly selected over the entire 15mm×15mm surface.

### AFM data analysis

The measurements were processed with in-house developed Matlab software.

### Lentivirus Construction and Generation

*E. coli* strains producing plasmids pEF.CMV.RFP (Addgene #17619, pEF.CMV.RFP was a gift from Linzhao Cheng), psPAX2 (Addgene #12260, psPAX2 was a gift from Didier Trono), and pMD2.G (Addgene #12259, pMD2.G was a gift from Didier Trono) were inoculated in LB Broth supplemented with ampicillin. Plasmid DNA was miniprepped (Qiagen). Human and mouse reference RNA were converted to cDNA as described above. Sequence for the gene of interest was PCR amplified from the reference cDNA using primers and cycles in **Supplementary Table 3**. Subsequent PCR product and pEF.CMV.RFP was digested with EcoRV according to the manufacturer’s protocol (NEB). These were then ligated together using T4 ligase (NEB) and transformed into STBL3 bacteria. Colonies were screened with PCR and positive clones were confirmed with sequencing yielding pLVmFut2 and pLVhFUT2.

### Intratracheal Elastase Administration and Pulmonary Function Tests

4.5U of elastase were intratracheally administered as previously described (15, 26). Briefly, mice were anesthetized with xylazine-ketamine mixture. The trachea was visualized, and tracheas were cannulated. 4.5U (50 uL) of elastase was administered and then mice were ventilated for 20 seconds. Mice were harvest 21 days following elastase administration. After 21 days mice were again anesthetized, and pulmonary function tests were performed with a FlexiVent(26). Lungs were then inflated with formalin and sent to Oncology Tissue Services (SKCCC, Baltimore MD) for paraffin embedding and mounting on slides. Mean linear intercept was quantified as previously described (37).

### Precision Cut Lung Slices (PCLS)

PCLS were prepared as previously described (38). PCLS were cut to be 250 microns thickness with a vibrating blade vibratome (Microm HM650V) and allowed to acclimate for 48 hours in DMEM F12 supplemented with 1% insulin-transferrin-selenium, and 1% antibiotic-antimycotic. Slice viability was assayed with Alamar Blue (ThermoFisher). Slices were transduced with fresh viral supernatant supplemented with 20 ug/mL of polybrene for 48 hours and then switched to fresh media for 24 hours before CS exposure. PCLS were exposed to CS or humidified air using the Vitrocell Systems GmbH smoking chamber with a previously described exposure protocol. PCLS were exposed to 8 cigarettes, with 1 cigarette every 8 minutes. After the last CS exposure, PCLS were returned to fresh media and incubated overnight. Following fixation, PCLS were processed by Oncology Tissue Services. MLI was quantified as above.

### Immunofluorescence

Deparaffinization, antigen retrieval, and immunofluorescence were performed as described previously (21). Slides were deparaffinized with xylenes followed by ethanol rehydration. Antigen retrieval was performed using Citrate Buffer (pH:6.0, ThermoFisher) for paraffin embedded tissues. Ki67 primary antibody (MA5-14520, ThermoFisher) and HOPX (11419, ProteinTech) were diluted 1:100 and SPC (518029, Santa Cruz Biotechnology (SCBT)) and CLDN4:AlexFluor488 (376643, SCBT) were diluted 1:50 and incubated overnight. AlexaFluor 555 and AlexaFluor 647 were diluted 1:200 and incubated for 1 hour. After secondary antibody incubation, the slides were incubated for 30 minutes in NucBlue™ Live ReadyProbes™ Reagent (ThermoFisher #R37605). After 30 minutes the slides were washed three times with PBS and mounted using the ProLong™ Glass Antifade Mountant (ThermoFisher #P36980). All immuofluorescent images were taken using the Johns Hopkins School of Medicine Microscope Facility Zeiss LSM700 Confocal.

### Cell Culture

Primary non-diseased human bronchial epithelial and COPD human bronchial epithelial cells were purchased from the Marsico Lung Institute (University of North Carolina Chapel Hill) expanded on collagen coated T75 flasks. Growth and differentiation conditions utilized were previously described (15, 16, 31, 38). A549 cells were a gift from Dr. Joseph Bressler (Johns Hopkins Bloomberg School of Public Health) and maintained in F12K media supplemented with 10% fetal bovine serum and 1% penicillin streptomycin. Cells were transduced with lentiviral supernatant for 48 hours supplemented with 20 ug/mL of polybrene. A549s that were transduced were then cell sorted at the Ross Flow Cytometery Core on a FACS Aria IIu Sorter as previously described (38) for RFP (and eGFP).

### Isolation of mice tracheal epithelial cells (mTECs)

Mouse tracheal epithelial cells were isolated as previously described (15, 38). Briefly, mice were euthanized by following carbon dioxide narcosis followed by cervical dislocation. The trachea were dissected out and added to 1X-Phosphate Buffer Saline (1X-PBS, ThermoFisher Scientific, New York, USA) supplemented with Penicillin-Streptomycin (ThermoFisher Scientific, NY, USA). Following incubation, the trachea were transferred to 0.15% Pronase solution and incubated overnight at 4°C. The solution was agitated and passed through a 70 µm cell strainer (Corning Life Sciences, MA, USA). Cells were pelleted from solution and resuspended in DMEM supplemented with FBS and Penicillin-Streptomycin. Cells were allowed to incubated in a flask for 4 hours to allow fibroblasts and mononuclear cells to attach. The remaining epithelial cells were transferred to a rat tail collagen I coated T75 flask and expanded in PneumaCult™-Ex Plus Basal Medium Supplemented with 10 mL PneumaCult™-Ex Plus 50X Supplement, 0.5 mL Hydrocortisone stock solution and 5 mL of 1% Penicillin-Streptomycin: StemCell Technologies Inc., Vancouver, Canada). At subconfluency cells were transferred to Transwell® (Corning) inserts and when confluent were allowed to differentiate at ALI for two weeks.

### Cigarette-smoke (CS) exposure to mTECs

The mTECs at 2 weeks ALI were exposed to either exposed to CS smoke or humidified air for 4 days as we previously described (15, 38). One CS exposure consisted of 2 cigarettes which burned for ∼ 8 minutes using the ISO puff regimen.

### Barrier Function Analysis

To determine monolayer integrity of the human and murine bronchial and tracheal epithelium at ALI, TEER was measured using epithelial voltohmeter (EVOM, World Precision Instruments Inc, FL, USA) with the STX2 electrodes as previously described (15). Values were corrected for fluid resistance and surface area. The paracellular permeability of the epithelium at ALI was determined using fluorescein isothiocyanate-dextran (FITC-Dextran) flux assay as described previously (15, 36).

## Acknowledgements

For creation of stable cell lines, cells were sorted at the JHU Ross Flow Cytometry Core. Fixed tissues were processed by Oncology Tissue Services at the Sidney Kimmel Comprehensive Cancer Center funded by grant P30 CA006973.

Research reported in this publication was supported by the National Heart, Lung, and Blood Institute (R01HL151107 and R01HL124099 to VKS), the Ludwig Family Department of Medicine Physician-Scientist Grant (VKS), and the Office of the Director of the National Institutes of Health under award number S10OD016374 (SC Kuo – JHU Microscope Facility).

---

**SF1:**
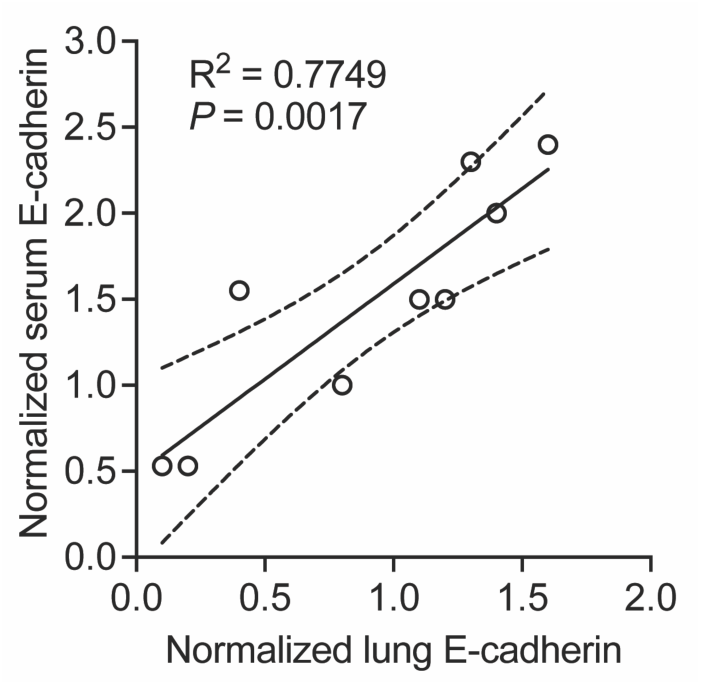
Serum levels of E-cadherin are highly correlated with lung levels of E-cadherin

**SF2:**
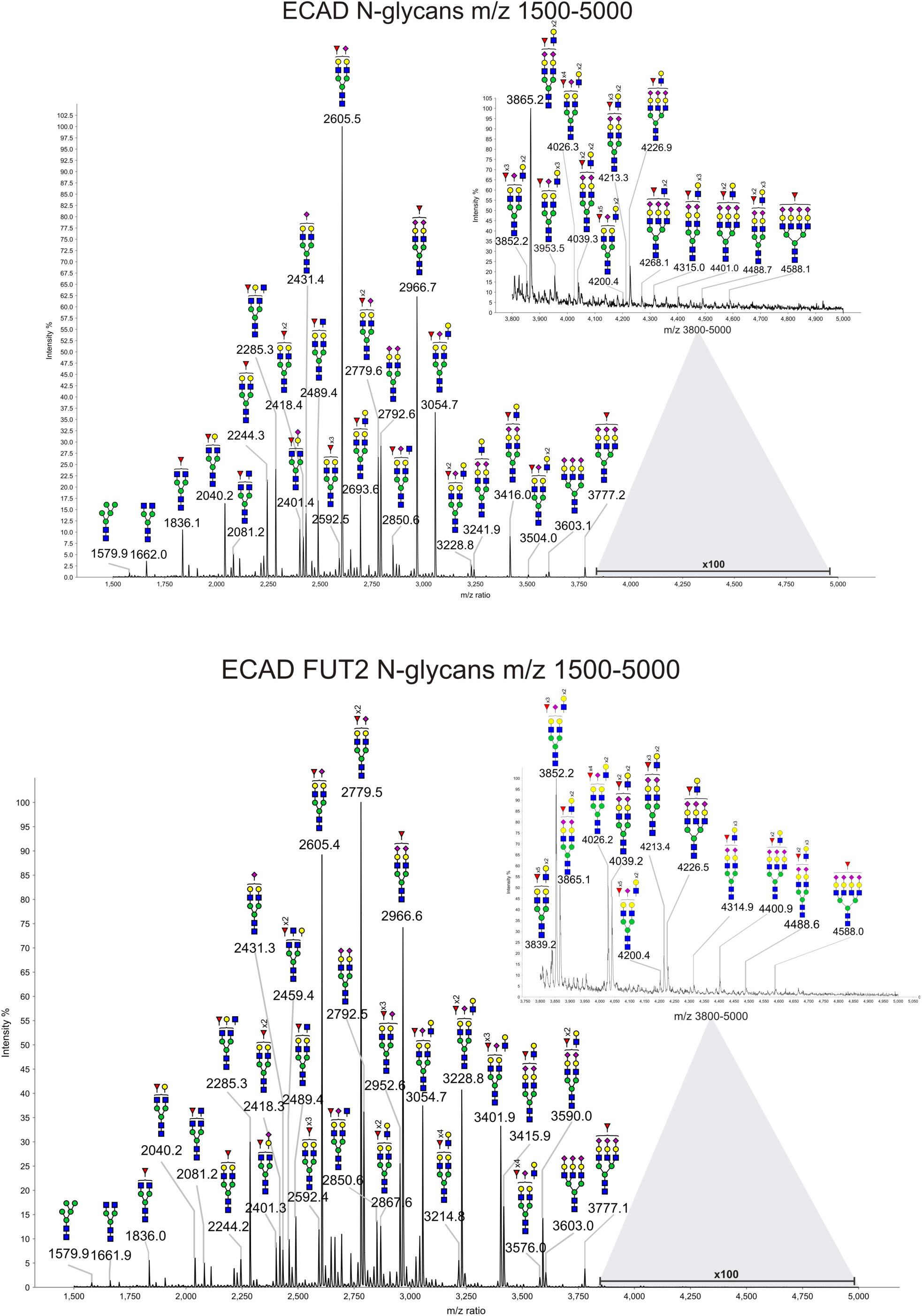
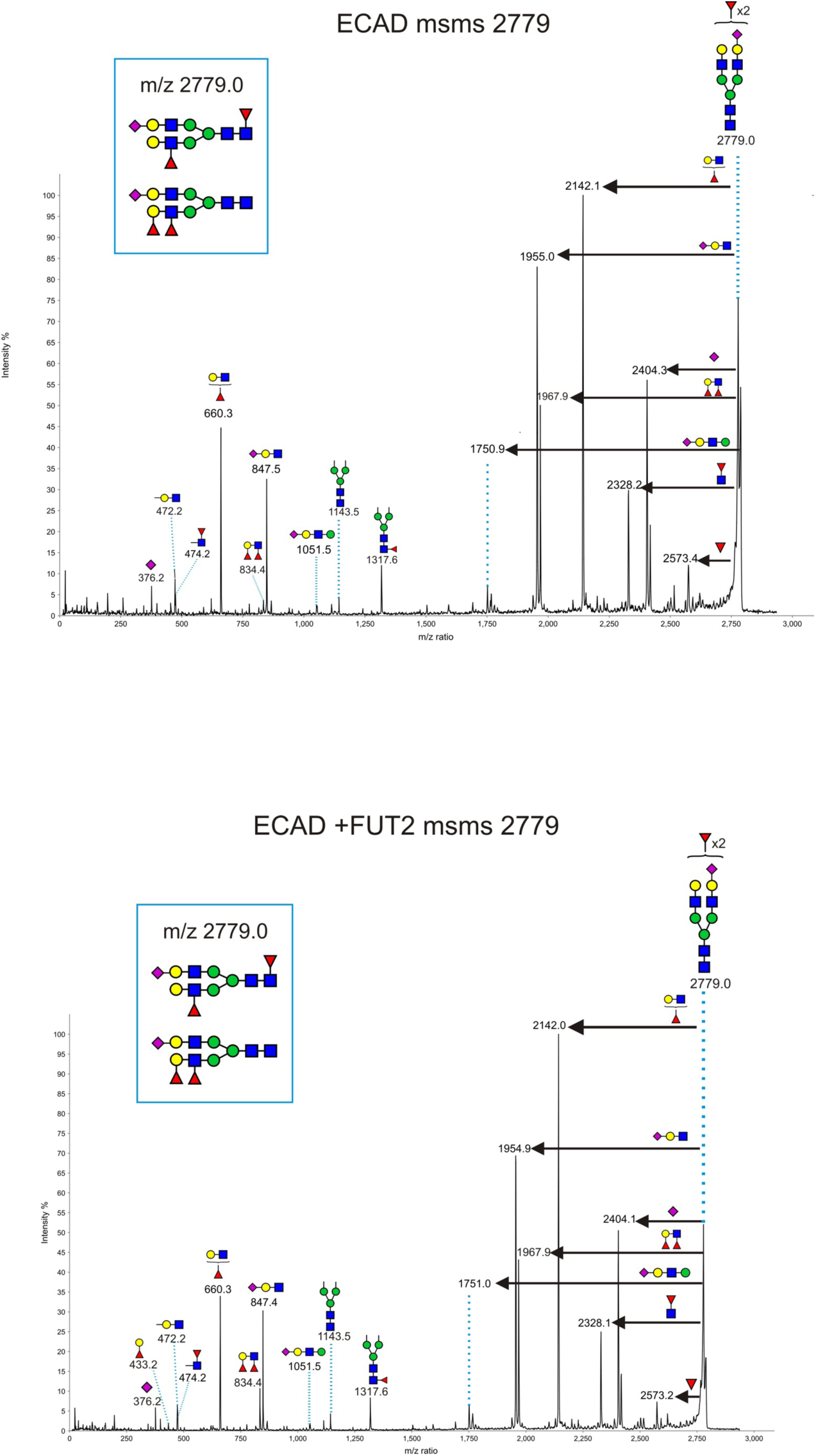
Glycan profiles of E-cadherin change with FUT2 overexpression

**SF3:**
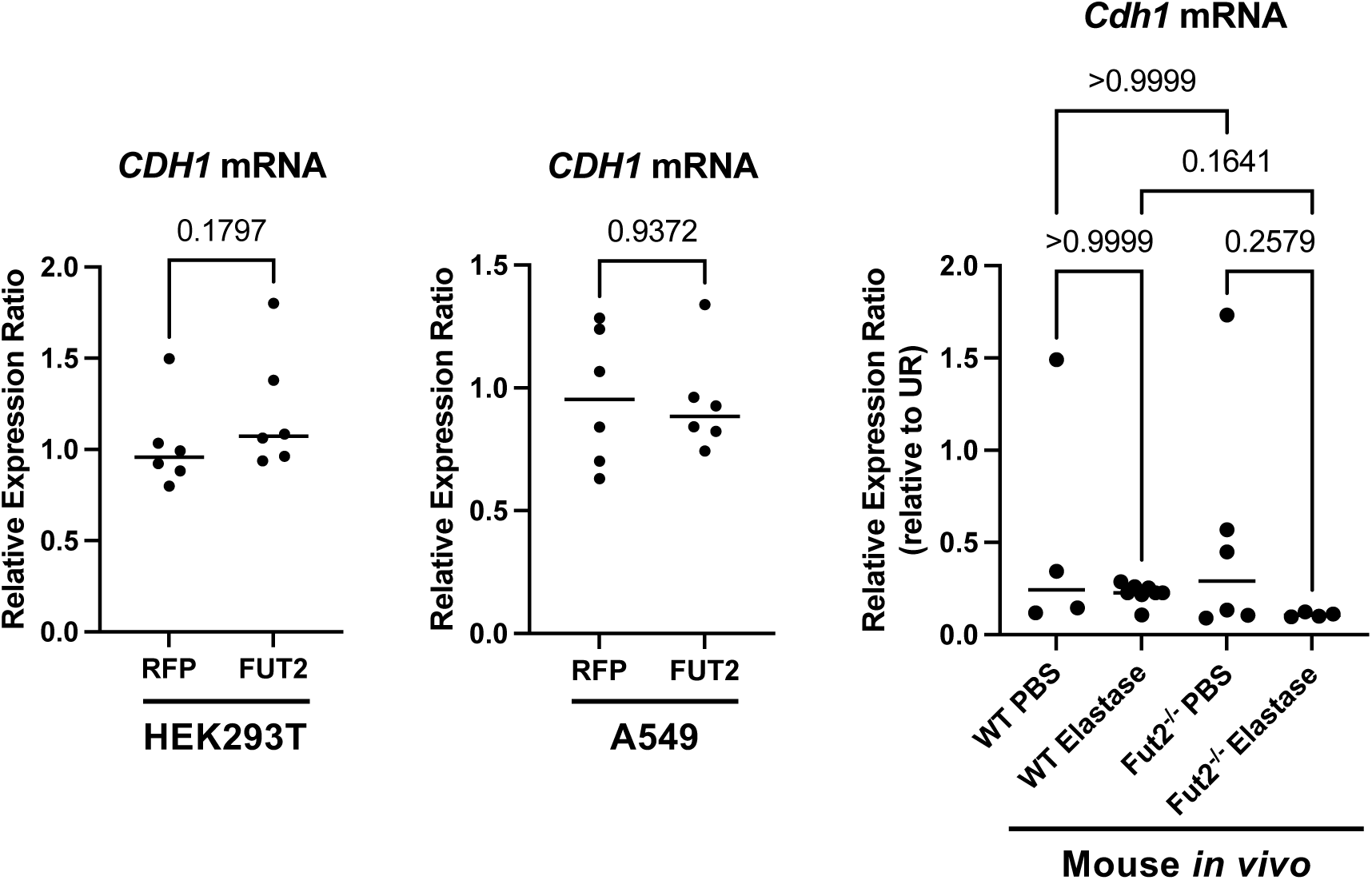
*CDH1* transcript levels are not dependent on *FUT2* expression.

**SF4:**
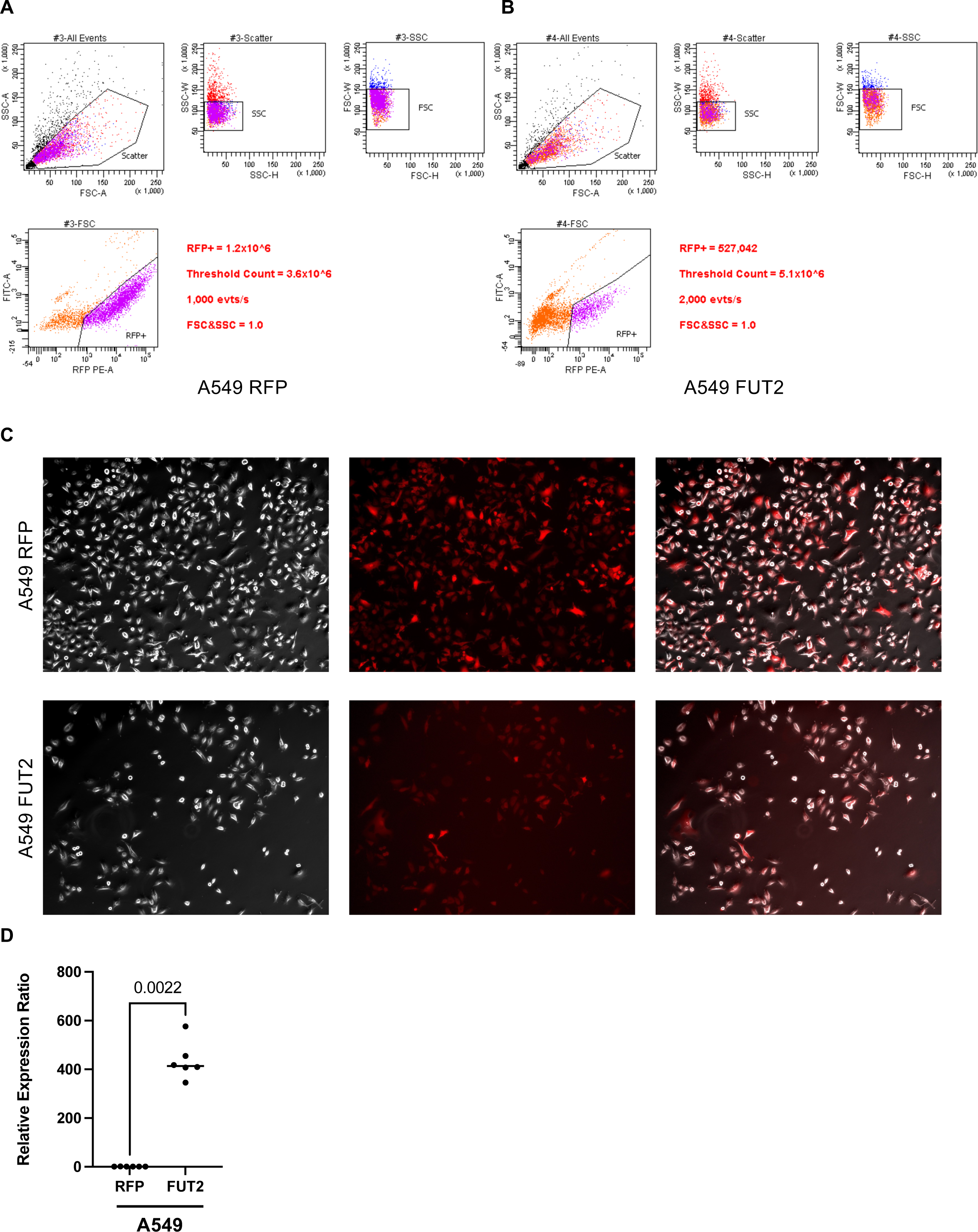
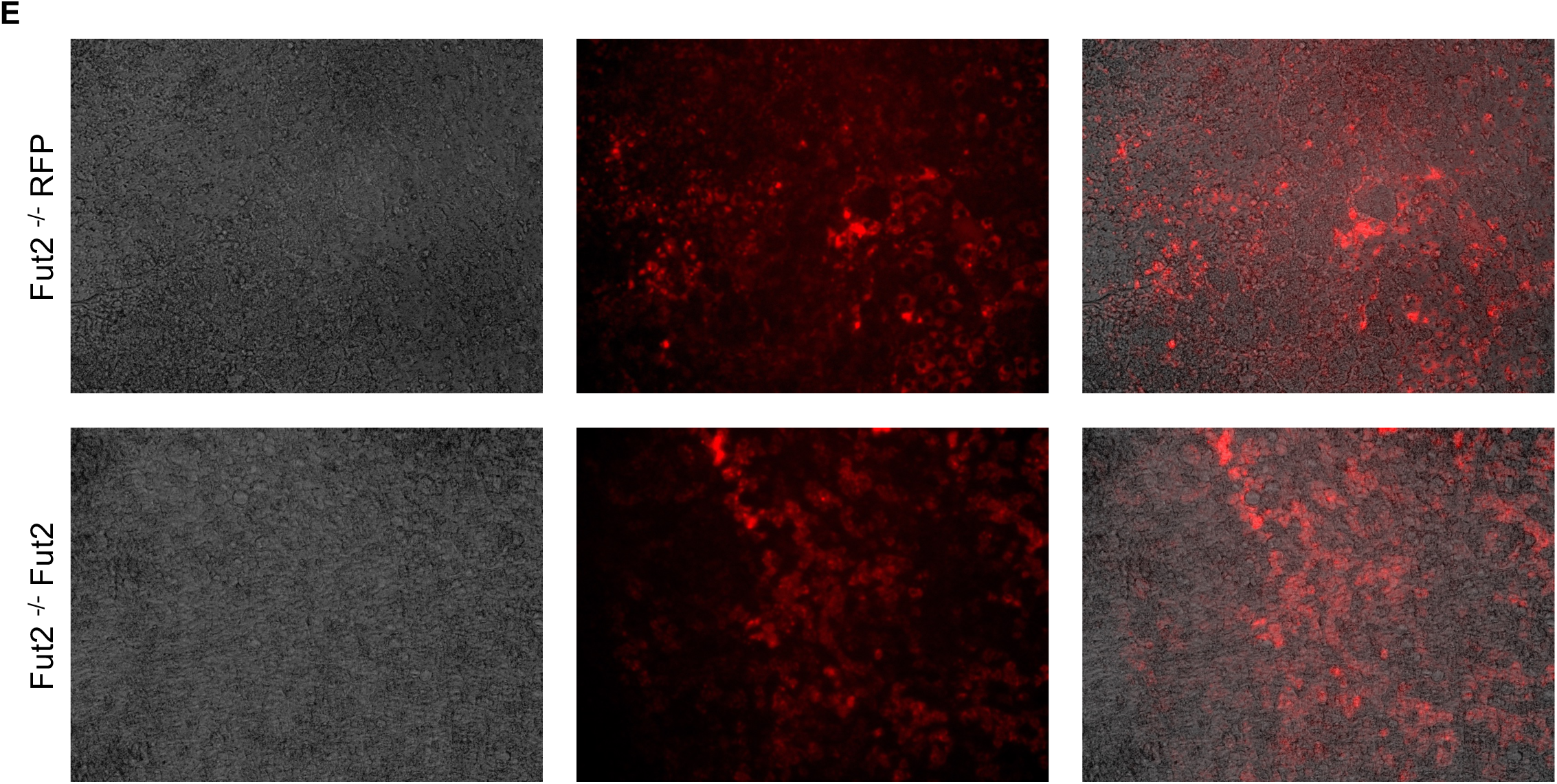
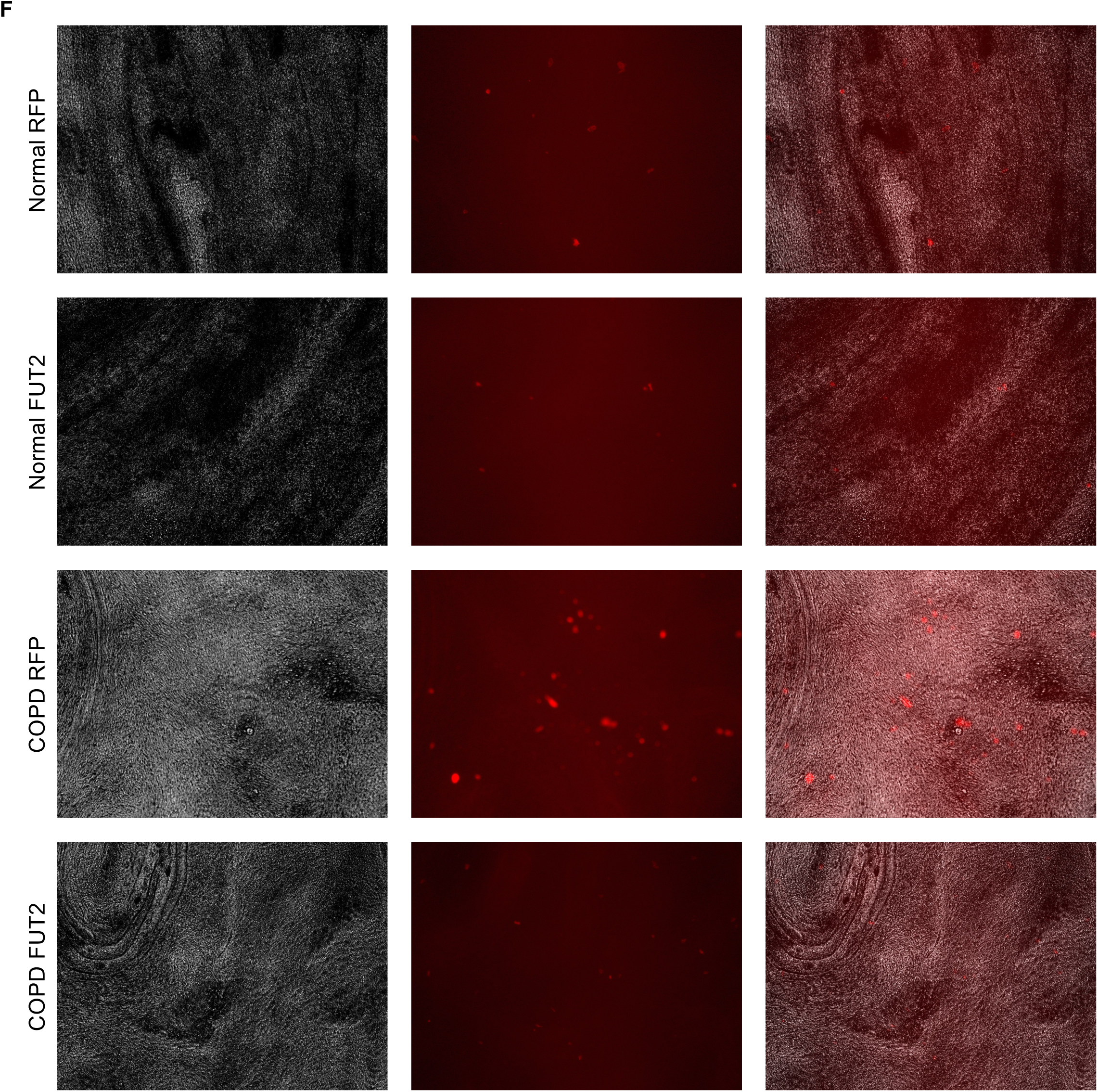
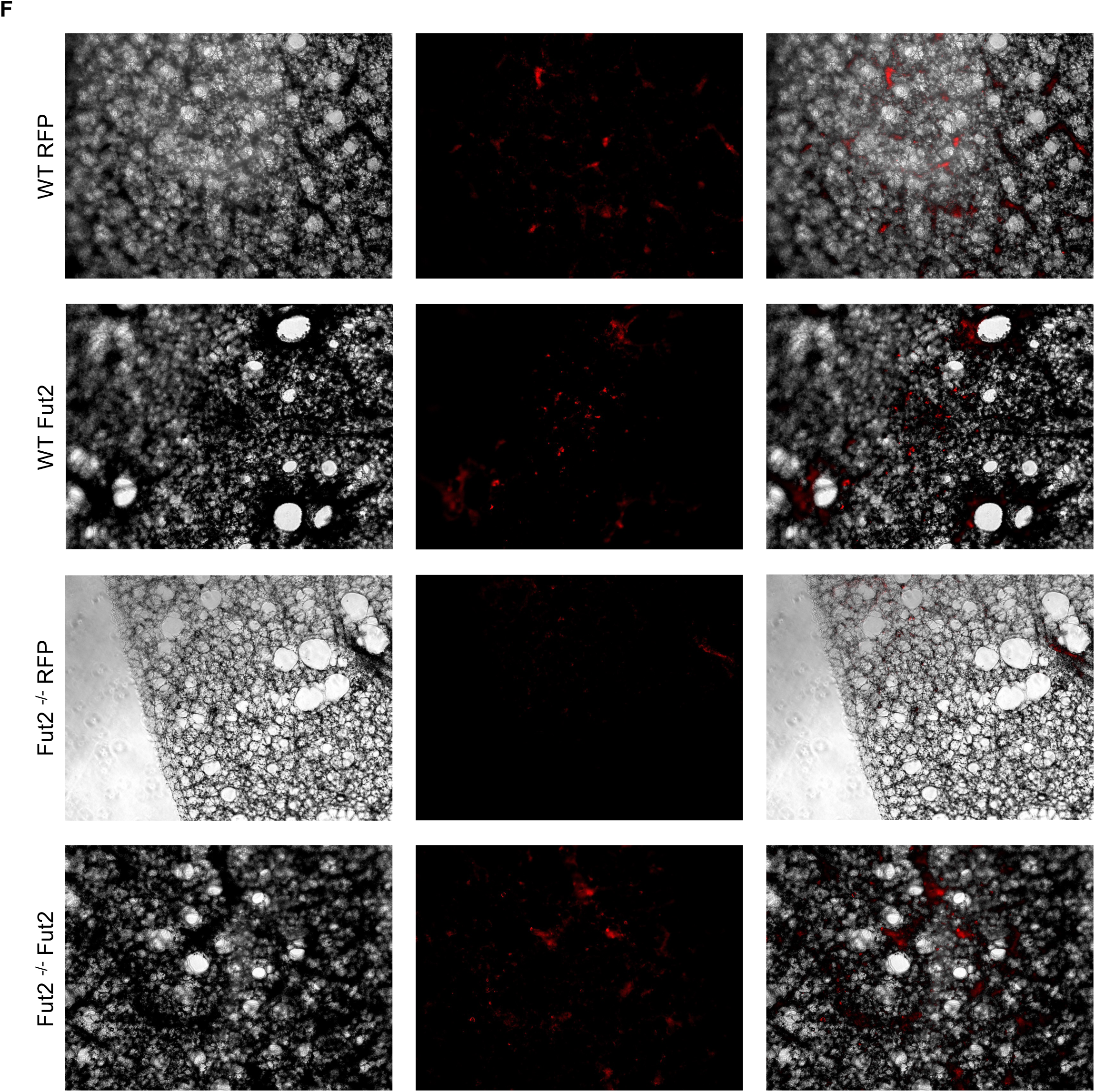
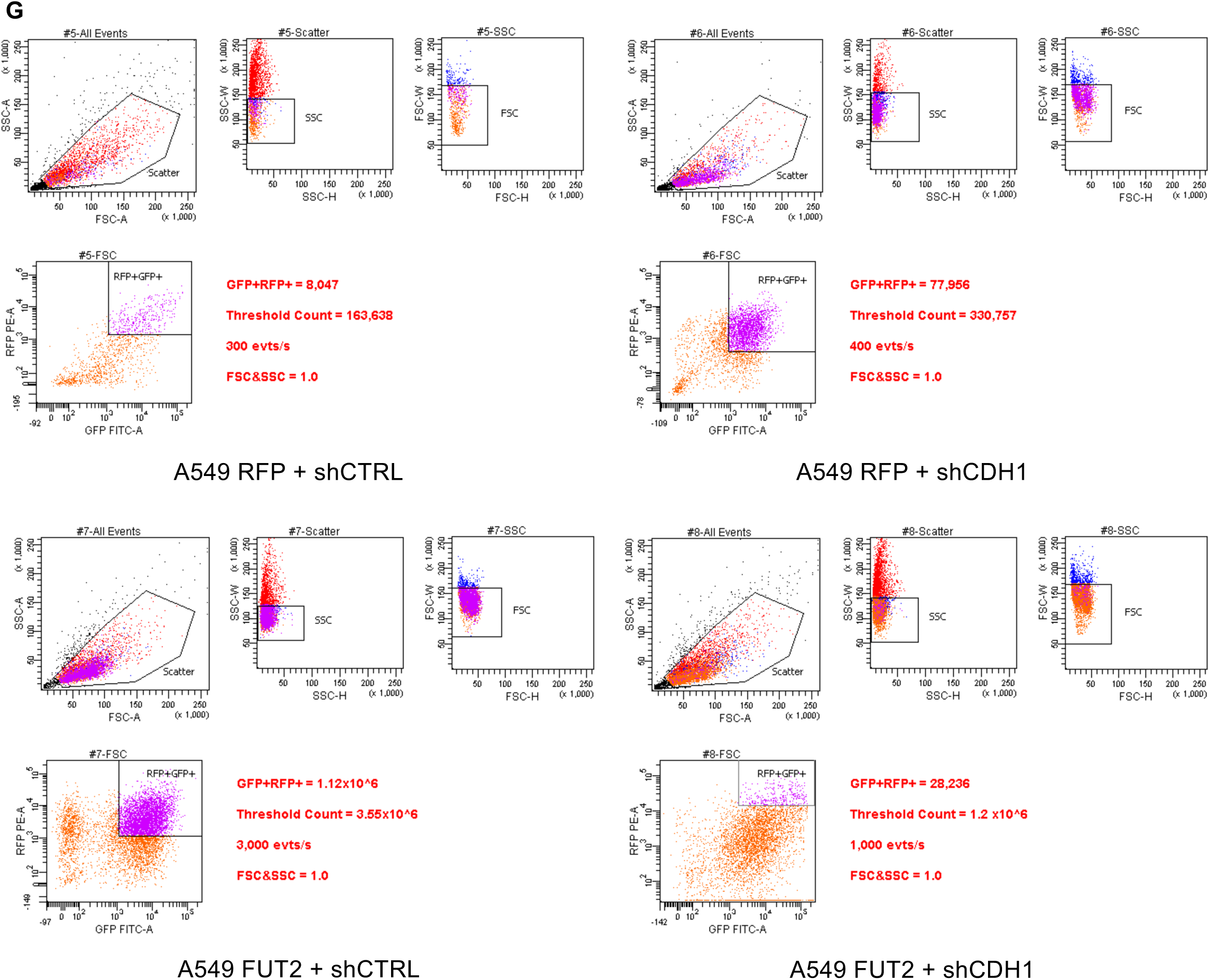
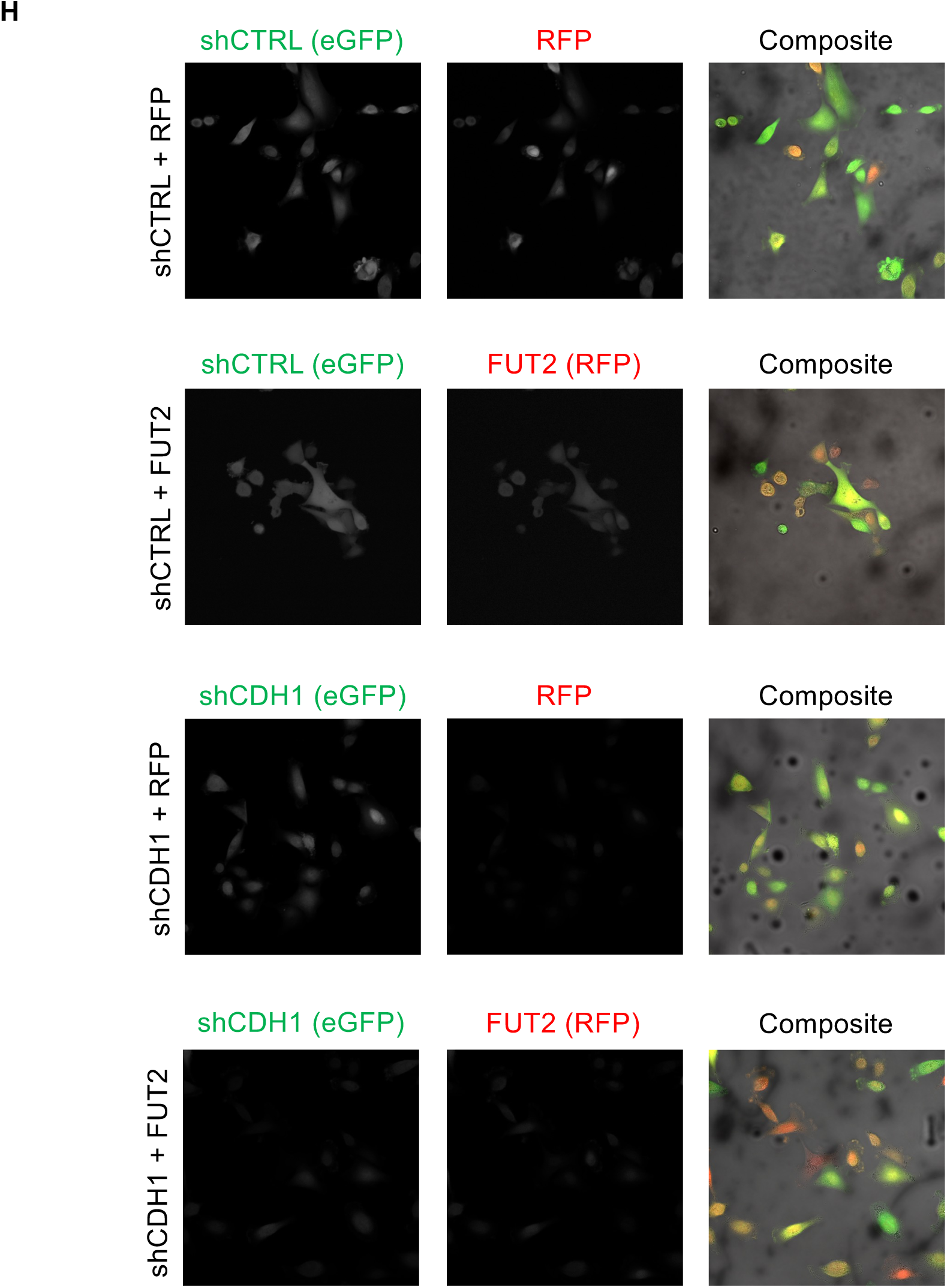
Transduction efficiency of RFP, FUT2, Fut2, shCTRL, shCDH1

**SF4:**
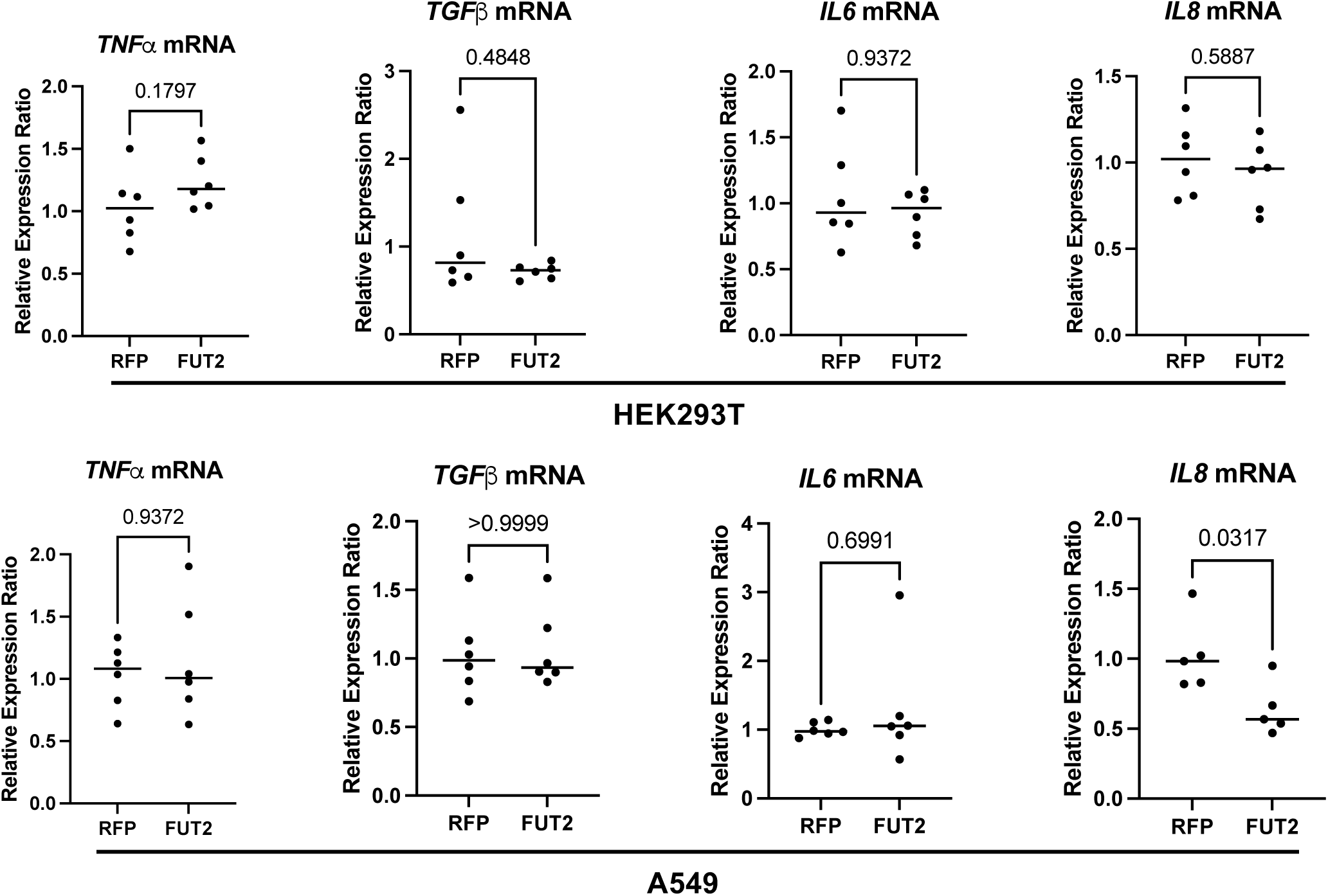
Markers of key inflammatory genes are not changed with varying FUT2 expression

**Supplementary Table 1:** Cohesion measurements of E-cadherin with and without FUT2 dependent fucosylation.

|  |  |  |
| --- | --- | --- |
| **E-cadherin without FUT2 fucosylation** |  |  |
| **Run** | **Adhesive Force (nN)** | **n** |
| 1 | 63.759 ± 2.8563 | 96 |
| 2 | 63.428 ± 3.5936 | 97 |
| 3 | 63.0181 ± 2.9007 | 101 |
| 4 | 63.4037 ± 2.9181 | 102 |
| 5 | 63.6881 ± 2.3771 | 101 |
| **E-cadherin with FUT2 fucosylation** |  |  |
| **Run** | **Adhesive Force (nN)** | **n** |
| 1 | 79.5199 ± 2.5149 | 105 |
| 2 | 80.2063 ± 3.5990 | 121 |
| 3 | 80.2640 ± 2.6532 | 117 |
| 4 | 79.6411 ± 3.0411 | 116 |
| 5 | 79.3105 ± 2.0907 | 107 |

**Supplementary Table 2:**
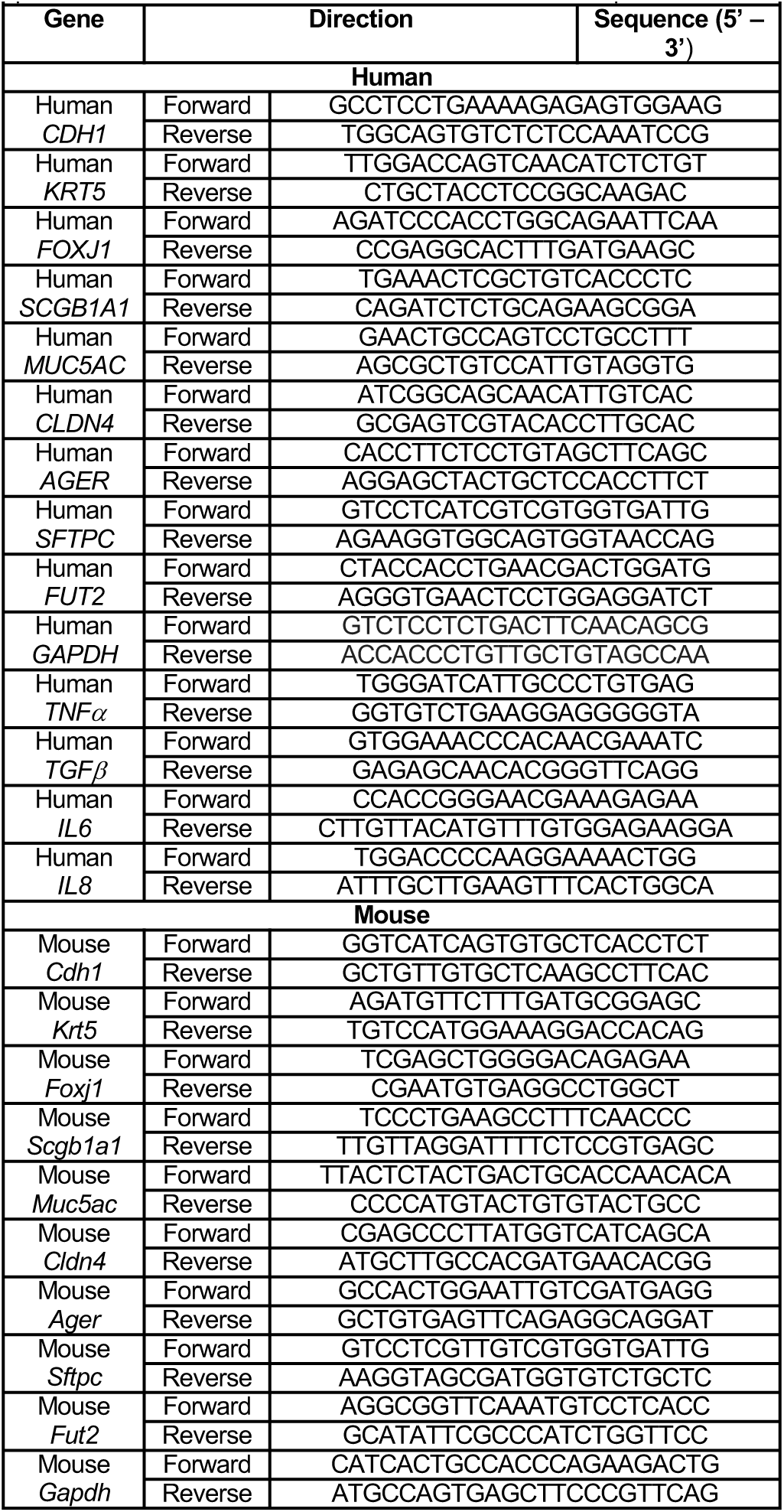
Quantitative polymerase chain reaction primers used to determine abundance of mRNA transcripts.

**Supplementary Table 3:**
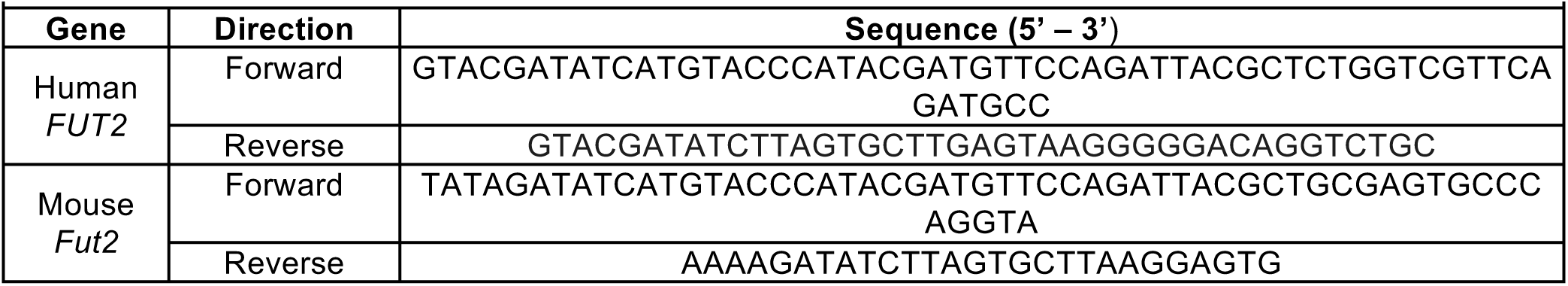
Primers used to amplify genes with polymerase chain reaction from genomic cDNA.

